# Real-time capture of σ^N^ transcription initiation intermediates reveals mechanism of ATPase-driven activation by limited unfolding

**DOI:** 10.1101/2025.02.07.637174

**Authors:** Andreas U. Mueller, Nina Molina, Seth A. Darst

**Affiliations:** Laboratory of Molecular Biophysics, The Rockefeller University, New York, NY, 10065 USA

## Abstract

Bacterial σ factors bind RNA polymerase (E) to form holoenzyme (Eσ), conferring promoter specificity to E and playing a key role in transcription bubble formation. σ^N^ is unique among σ factors in its structure and functional mechanism, requiring activation by specialized AAA+ ATPases. Eσ^N^ forms an inactive promoter complex where the N-terminal σ^N^ region I (σ^N^-RI) threads through a small DNA bubble. On the opposite side of the DNA, the ATPase engages σ^N^-RI within the pore of its hexameric ring. Here, we perform kinetics-guided structural analysis of *de novo* formed Eσ^N^ initiation complexes and engineer a biochemical assay to measure ATPase-mediated σ^N^-RI translocation during promoter melting. We show that the ATPase exerts mechanical action to translocate about 30 residues of σ^N^-RI through the DNA bubble, disrupting inhibitory structures of σ^N^ to allow full transcription bubble formation. A local charge switch of σ^N^-RI from positive to negative may help facilitate disengagement of the otherwise processive ATPase, allowing subsequent σ^N^ disentanglement from the DNA bubble.

## Main Text

Specific promoter recognition during bacterial transcription initiation requires the association of a σ factor with the RNA polymerase (RNAP) core enzyme (α_2_ββ’ω; E) to form the holoenzyme (Eσ) ^1^. Multiple alternative σ factors compete for E, enabling the cell to switch transcriptional programs. Alternative σ factors are almost all homologous to the housekeeping *Escherichia coli* (*Eco*) σ^70^ factor ^2^, but nearly all bacterial clades except Actinobacteria harbor a structurally and evolutionarily unrelated σ factor, σ^N^ (also known as RpoN or σ^54^) ^3,4^. σ^N^ mediates diverse transcriptional pathways, most notably the expression of virulence factors in many human bacterial pathogens including *Pseudomonas aeruginosa*, *Vibrio cholerae*, the Lyme disease agent *Borrelia burgdorferi*, *Chlamydia trachomatis*, and *Helicobacter pylori* ^5–9^, and the control of nitrogen metabolism in plant symbionts such as *Rhizobium sp.* ^10^.

Unlike factors of the σ^70^ family, transcription initiation by Eσ^N^ proceeds through a stable, inactive promoter complex that requires ATP-dependent post-recruitment activation mediated by a bacterial enhancer-binding protein (bEBP) ^3,4^. The bEBPs, which belong to the large superfamily of AAA+ proteins (ATPases associated with diverse cellular activities), activate transcription through an unknown mechanism. All known bEBPs form hexameric rings and consist of a central AAA+ domain, with variable presence of an N-terminal regulatory domain and/or a C-terminal DNA binding domain enabling recognition of promoter-distal enhancer sequence motifs ^11^. Full DNA melting and progression to the transcription-competent ‘open promoter complex’ (RPo) are prohibited by an interaction of σ^N^ N-terminal region I (RI) with the extra-long helix (ELH) (Extended Data Fig. 1A) ^12–15^. At promoters, Eσ^N^ recognizes conserved sequence elements located 12 and 24 base pairs upstream (the −12 and −24 elements) of the transcription start site (TSS; position +1) via the σ^N^ ELH helix-turn-helix (HTH) domain and RpoN domain, respectively ^16–18^, forming a stable early melted intermediate (RPem) in which two base pairs (at −12 and −11) are melted ^19^. In RPem, an N-terminal segment of σ^N^-RI threads through the DNA bubble ^20,21^. The bEBP engages σ^N^-RI on the opposite side of the DNA bubble with highly conserved GAFTGA loops in the AAA+ domain ^20,22^.

Recent structural work revealed the interaction of σ^N^-RI inside the pore of the hexameric ring ^20^. The structure is trapped with ADP aluminum fluoride (ADP-AlFx) but leads to an attractive hypothesis: the ATP hydrolysis cycle of the AAA+ bEBP pulls σ^N^-RI through its pore, mechanically unfolding σ^N^-RI and disrupting the σ^N^-RI:ELH interactions. The mechanical disruption of the σ^N^-RI:ELH interaction could explain how bEBP action allows RPo formation to proceed, but this mechanistic hypothesis also raises crucial questions: How many rounds of ATP hydrolysis are required to disrupt the σ^N^-RI:ELH interactions? Most AAA+ translocases are highly processive ^23^; does the bEBP thread the entire length of the σ^N^ polypeptide (*Eco* σ^N^ is 477 residues in length) through the DNA bubble? Alternatively, the bEBP could disengage from σ^N^ after RPo formation, but what is the signal for disengagement?

To address these questions, we perform a kinetics-guided structural analysis of *de novo* formed Eσ^N^ initiation complexes using cryo-electron microscopy (cryo-EM) and engineer a biochemical assay to measure bEBP-mediated σ^N^-RI translocation in initiation complexes. We show that the bEBP translocates about 30 residues of σ^N^-RI through the DNA bubble, converting ATP hydrolysis into mechanical force to disrupt the σ^N^-RI:ELH inhibitory interface to allow full transcription bubble formation. Our data suggests that a switch of local net charges in σ^N^-RI from positive to negative may facilitate disengagement of the otherwise processive bEBP, permitting σ^N^ disentanglement from the DNA bubble during later steps.

## Results

### Kinetics-informed capture of actively initiating Eσ^N^-bEBP complexes by cryo-EM

To capture and structurally visualize intermediates during bEBP action, we first determined the kinetics of RPo formation. We employed a fluorescence assay where a Cy3 fluorophore attached to the +2 position experiences protein-induced fluorescence enhancement (PIFE) ^24^ upon RPo formation (Fig. 1A) ^25^. Our transcription system comprised *Eco* Eσ^N^, a consensus σ^N^ promoter (the *Aquifex aeolicus* dhsU promoter; ^16,26^), and a constitutively active variant of the *A. aeolicus* (*Aae*) bEBP NtrC1 (C1; N-terminal regulatory domain and C-terminal enhancer DNA-binding domain removed) ^27^. To enable correct positioning of the fluorescent label, we determined the TSS for both the wild-type (+2A) and mutant (+2T) promoters (Cy3 is attached via a T base) under our experimental conditions (Supplementary Table 1). A Cy3-labelled dhsU promoter fragment was synthesized (dhsU+2T-Cy3; Supplementary Table 2) and used in a stopped flow assay to determine the kinetics of RPo formation. The fluorescence signal reached a plateau after about 5 min, indicating RPo formation was complete (Fig. 1B). Kinetic analysis of the data yielded a maximum apparent rate constant for RPo formation of about 0.009 s^−1^ (Extended Data Fgigs. 1B-D). Accounting for the difference in reaction temperature, these results are consistent with previously reported data ^28^.

**Fig. 1:**
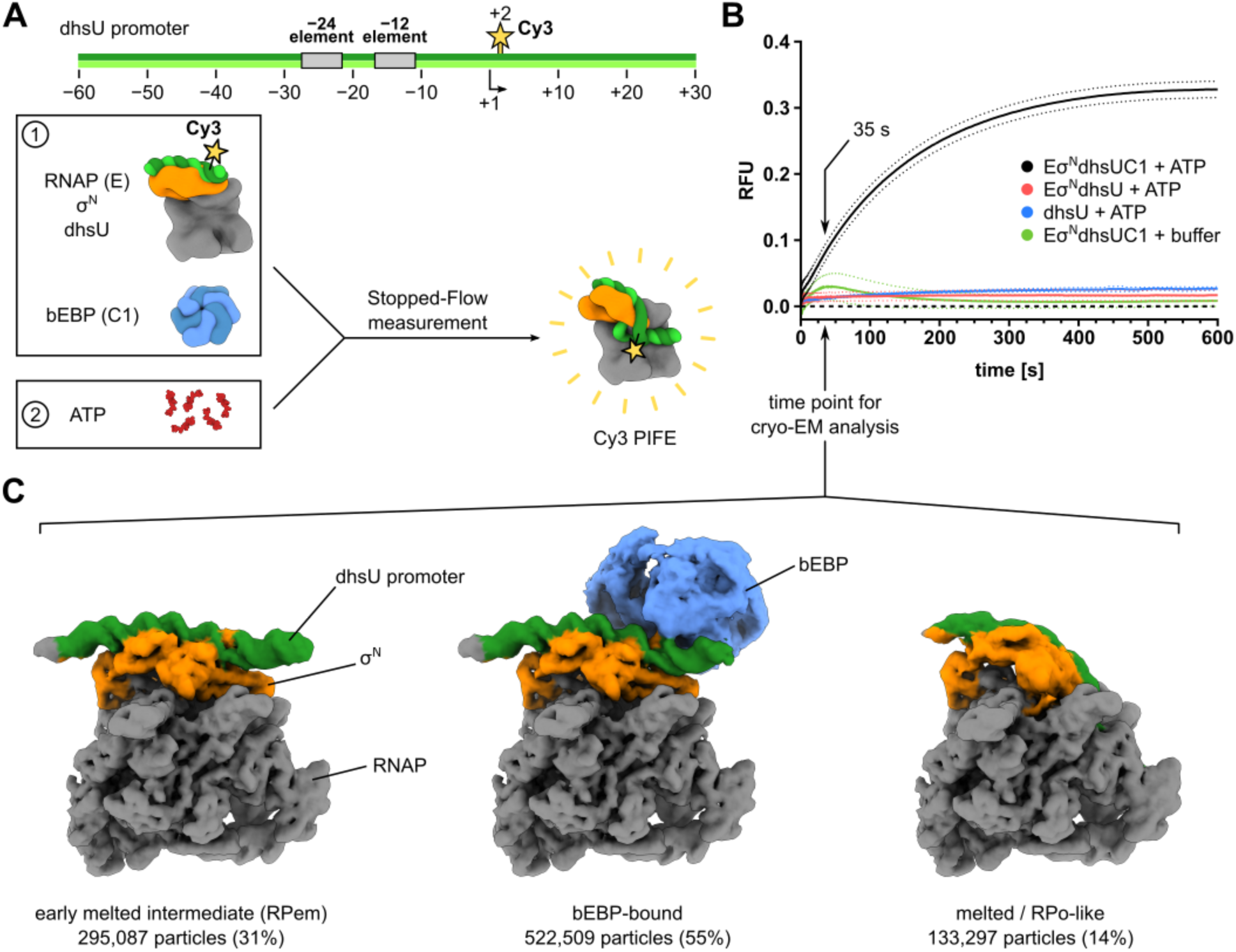
RPo formation kinetics of Eσ^N^ informs cryo-EM analysis. (A) Stopped-flow measurement employs a linear, Cy3-labeled (yellow) σ^N^ promoter DNA (*Aae* dhsU). The fluorophore is subject to protein-induced fluorescent enhancement (PIFE) when RPo forms. For the experiment, an early-melted intermediate (box 1; E+σ^N^+dhsU+C1) is pre-assembled from core RNAP (E, grey), σ^N^ (orange), and Cy3-labeled dhsU (green) in presence of bEBP (C1; blue). ATP is prepared in buffer (box 2). Mixing the contents of boxes 1 and 2 will start the reaction and protein-induced fluorescence enhancement (PIFE) can be observed. (B) RPo formation-dependent fluorescence saturates at around 8-10 minutes reaction time and is dependent on the presence of all components (black). Mixing ATP with complexes without C1 (red), ATP with DNA alone (blue), or the E+σ^N^+dhsU+C1 with buffer (no ATP, green) results in only background signal. Dotted lines indicate error bands of one standard deviation. Each trace is an average of 5-7 reactions. The arrow indicates the time point used for cryo-EM sample preparation. (C) Cryo-EM analysis of samples prepared after 35 seconds reaction time yields three major populations: early-melted intermediates (RPem, 31% of particles), bEBP-bound complexes (55% of particles), and complexes with a melted promoter DNA (RPo-like, 14%). Maps from the 3D classification (see main text and methods for details) were colored according to core RNAP (grey), σ^N^ (orange), dhsU (green), and bEBP (blue).

The population of melting intermediates should be greatest around the initial rise of the signal, which occurred around 20-50 seconds (Fig. 1B). Accordingly, cryo-EM samples were prepared by adding ATP to RPem + C1 and plunged into liquid ethane after 35 seconds reaction time (Fig. 1B). Data processing revealed a class of bEBP-bound complexes (ca 55%), a class of RPem complexes without bEBP (31%), and a class of RPo-like complexes (no bEBP bound, DNA melted and inserted into the active site cleft of RNAP; ca 14%) (Fig. 1C; Extended Data Fig. 2A). The fraction of RPo-like complexes (14%) roughly corresponds with the fraction of maximum signal obtained during the stopped flow assay at 35 seconds (25%), establishing consistency between the different experimental approaches (Figs. 1B-C) (note that direct correspondence between the two assays is not expected since the fluorescence signal in the stopped flow assay is not a linear function of RPo abundance).

### Structures of Eσ^N^-bEBP initiation complexes reveal bEBP-mediated translocation and unfolding of σ^N^-RI

The bEBP-bound class displayed a large variability of the bEBP position around the axis along the σ^N^-bound DNA. Steps of focused classifications and local alignments (see Materials and Methods) revealed “open” and “closed” bEBP ring states defined by the presence or absence of a distinct gap in the ring, respectively (Extended Data Fig. 2A). For each ring state, maps encompassing the whole complex were obtained by merging the bEBP-focused map with a map reconstructing RNAP (i.e., without masks) and a σ^N^-focused map. Nominal FSC resolution estimates were similar across both ring states: 3.0-3.1 Å for the bEBP and σ^N^ maps, and 2.7 Å for the RNAP maps, with homogeneous distributions of angular views (Table 1; Extended Data Figs. 3 and 4).

**Table 1:** Cryo-EM map and model statistics. Calculation of model statistics was performed using MolProbity.

| Complex | RPi1<sup>open</sup> (open ring) | RPi1<sup>closed</sup> (closed ring) | RII | RPO | RPO+2A |
| --- | --- | --- | --- | --- | --- |
| PDB ID | 9MSE | 9MSF | 9MSG | 9MSH | 9MSJ |
| EMDB ID | EMD-48586<br>EMD-48580<br>EMD-48581<br>EMD-48582 | EMD-48587<br>EMD-48583<br>EMD-48584<br>EMD-48585 | EMD-48588 | EMD-48589<br>EMD-48576<br>EMD-48577 | EMD-48590<br>EMD-48578<br>EMD-48579 |
| Data collection |  |  |  |  |  |
| Camera | Gatan K3 |  |  |  |  |
| Dose rate (e<sup>-</sup>/Å<sup>2</sup>/s) | 30 |  |  |  |  |
| Electron exposure (e<sup>-</sup>/Å<sup>2</sup>) | 42 |  |  |  |  |
| Defocus range (μm) | -0.8 to -2.2 |  |  |  |  |
| Pixel size (Å) | 0.86 |  |  |  |  |
| Energy filter slit width (eV) | 20 |  |  |  |  |
| Cs (mm) | 0.001 |  |  |  |  |
| Images collected | 17,199 |  |  |  |  |
| Symmetry imposed | C1 |  |  |  |  |
| Initial particles | 2,798,596 |  |  |  |  |
| Consensus final particles | 1,060,259 |  |  |  |  |
| Consensus resolution (Å)<sup>a</sup> | 2.3 |  |  |  |  |
| Processing statistics |  |  |  |  |  |
| Class particles | 118,561 | 260,106 | 95,920 | 45,631 | 15,649 |
| bEBP map resolution (Å)<sup>a</sup> | 3.1 Å | 3.0 Å | n/a | n/a | n/a |
| Sharpening b-factor | -85.4 | -95.2 | n/a | n/a | n/a |
| RNAP map resolution (Å)<sup>a</sup> | 2.7 Å | 2.6 Å | 2.7 Å | 2.8 Å | 3.1 Å |
| Sharpening b-factor | -61.6 | -68.1 | -60.4 | -54.1 | -39.4 |
| σ<sup>N</sup> map resolution (Å)<sup>a</sup> | 3.1 Å | 3.0 Å | n/a | 3.2 Å | 3.6 Å |
| Sharpening b-factor | -74.2 | -80.3 | n/a | -67.2 | -55.5 |
| Refinement |  |  |  |  |  |
| Initial models used | 8F1K<br>4LZZ | 8F1K<br>4LZZ | 8F1K<br>4LZZ | 8F1K<br>6GH5 | 8F1K<br>6GH5 |
| Non-hydrogen atoms | 43,810 | 44,073 | 42,869 | 31,061 | 31,102 |
| Protein residues | 5,200 | 5,201 | 5,235 | 3,664 | 3,687 |
| Nucleotides | 92 | 92 | 68 | 98 | 104 |
| Water molecules | 703 | 986 | 29 | 348 | 54 |
| Ligands | 12 | 11 | 11 | 4 | 5 |
| RMS (bonds) | 0.003 | 0.002 | 0.002 | 0.003 | 0.003 |
| RMS (angles) | 0.458 | 0.446 | 0.459 | 0.515 | 0.505 |
| Ramachandran favored | 97.75 % | 98.18 % | 97.73 % | 97.20 % | 98.42 % |
| Ramachandran allowed | 2.23 % | 1.82 % | 2.27 % | 2.80 % | 1.58 % |
| Ramachandran outliers | 0.02 % | 0.00 % | 0.00 % | 0.00 % | 0.00 % |
| Rotamer outliers | 1.10 % | 1.12 % | 1.56 % | 2.74 % | 2.44 % |
| Clashscore | 3.36 | 3.62 | 4.72 | 4.62 | 4.64 |
| MolProbity score | 1.21 | 1.19 | 1.45 | 1.72 | 1.53 |

<sup>a</sup> FSC threshold 0.143

The structures of both ring states share several common features: Density for the dhsU promoter fragment is clearly resolved from about −35 to −2 (Fig. 2A). As in the RPem state (without bEBP) ^20,21^, σ^N^-RI threads between the opened DNA strands in the structures of both ring states (Figs. 2A and 2B). After σ^N^-RI threads through the opened DNA strands, it engages with the pore of the bEBP hexamer on the opposite side of the DNA (Figs. 2A, 2C-right panel; Extended Data Fig. 5) as previously seen in a bEBP-bound RPem trapped in the presence of ADP-AlF_x_ (RPem-bEBP^ADP-AlFx^; PDB 7QV9; Fig. 2C-left panel) ^20^. In RPem-bEBP^ADP-AlFx^, σ^N^-RI residues between 1-11 are captured in the pore ^20^ while our Eσ^N^-bEBP open and closed ring states (in the presence of ATP) captured σ^N^-RI residues between 1-13 – a two-residue register shift deeper into the pore (Fig. 2C; Extended Data Fig. 5B). The two-residue register shift indicates that the open and closed ring states captured in the presence of ATP underwent one translocation step, consistent with the two amino acid step size of protein translocating AAA+ ATPases ^23^. Therefore, we refer to the open and closed ring states as RPi1^open^ and RPi1^closed^ to indicate that they represent a translocation intermediate after one step of ATP hydrolysis.

**Fig. 2:**
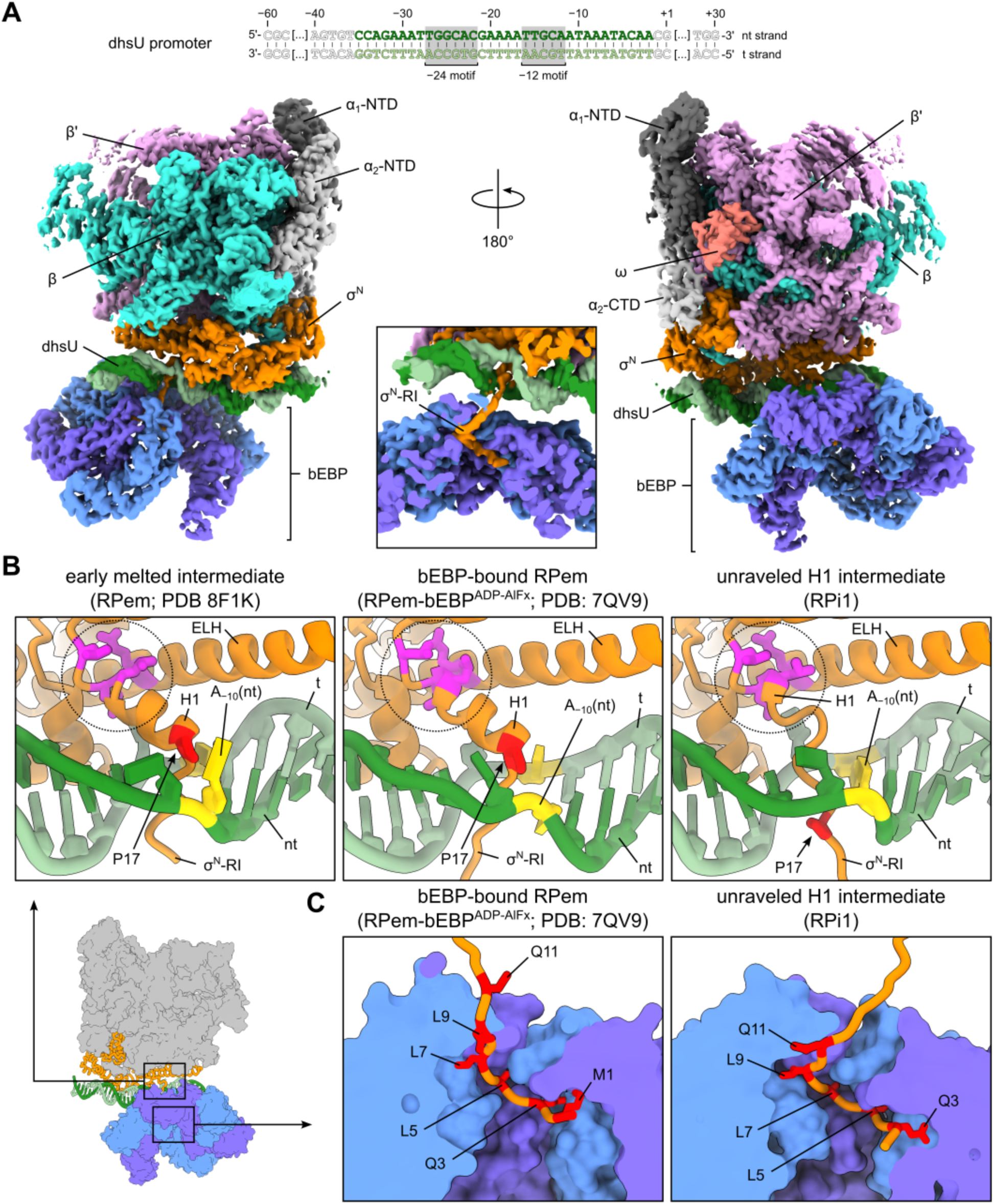
Cryo-EM structures capture translocation and remodelling of σ^N^-RI by the bEBP. (A) Overview of the merged, unsharpened cryo-EM map of the “open” ring state. Map density is colored according to the location of subunits. The extent of visible map density for the DNA duplex is indicated by the colored letters at the top. (B) Helix 1 (H1) of σ^N^-RI exhibits three turns in RPem (left) and RPem-bEBP^ADP-AlFx^ (middle), but a full turn is unfolded in the RPi1 intermediate (right). Residue P17 (red) located at the N-terminal end of H1 in RPem serves as visual indicator of helix changes. Regulatory interactions between σ^N^-RI and the ELH remain intact (dotted circle; residues in magenta are mutation sites for bypass variants). (C) Comparison of the σ^N^-RI:bEBP interaction between RPem-bEBP^ADP-AlFx^ (left) and RPi1 (right), illustrating the translocation of σ^N^-RI by two residues. Every second residue in σ^N^-RI starting with M1 up to Q11 is highlighted in red. M1 was not modeled for the open and closed ring structures.

In both RPem (PDB 8F1K) ^21^ and RPem-bEBP^ADP-Alx^ (PDB 7QV9) ^20^, Pro17 marks the N-terminal end of σ^N^ helix 1 (H1) and sits on the RNAP-side of the small DNA bubble accommodating the threaded σ^N^-RI (Fig. 2B-left and middle panels). By contrast, the bEBP-mediated translocation of σ^N^-RI N-terminal end in the RPi1 structures partly unravels H1, pulling Pro17 more than 9 Å through to the bEBP-side of the DNA bubble (Fig. 2B-right panel). The A-T base pair at the −10 position reforms yielding only a one nucleotide bubble consistent with the reduced spatial requirements for a partially folded H1.

All known ‘bypass’ mutants of σ^N^ – which allow RPo formation and transcription initiation without bEBP intervention ^12,14,15,29–31^ – participate in a network of conserved interactions that stabilize the interaction between σ^N^-RI and the σ^N^-ELH-HTH domain (Fig. 2B) ^16^. Thus, this intra-protein interface is crucial for proper regulation of σ^N^ regulons by preventing unregulated transcription. This regulatory interface includes the C-terminal end of σ^N^-H1 but remains intact in the RPi1 structures despite the unraveling of the σ^N^-H1 N-terminal end (Fig. 2B-right panel). Further bEBP-mediated translocation of σ^N^-RI would presumably unravel σ^N^-H1 completely and disrupt the regulatory interface, allowing RPo formation to proceed.

### bEBP-mediated unfolding and threading of σ^N^-RI halts at a defined position

To observe and quantify the extent of the bEBP threading and unfolding activity indicated by the structural data, we engineered a proteolytic assay to measure the extrusion of σ^N^-RI during RPo formation. In this assay, the bEBP was engineered to interact with a compartmentalized protease, where it could translocate σ^N^-RI into the proteolytic chamber for degradation, providing a readout of the extent of σ^N^-RI extrusion (Fig. 3A). We designed our constructs after the mycobacterial proteasome and its regulator, the mycobacterial proteasome AAA+ ATPase (Mpa), which features a defined ATPase-proteasome interaction motif (GQYL) and well-characterized interaction to the proteasome core ^32^.

**Fig. 3:**
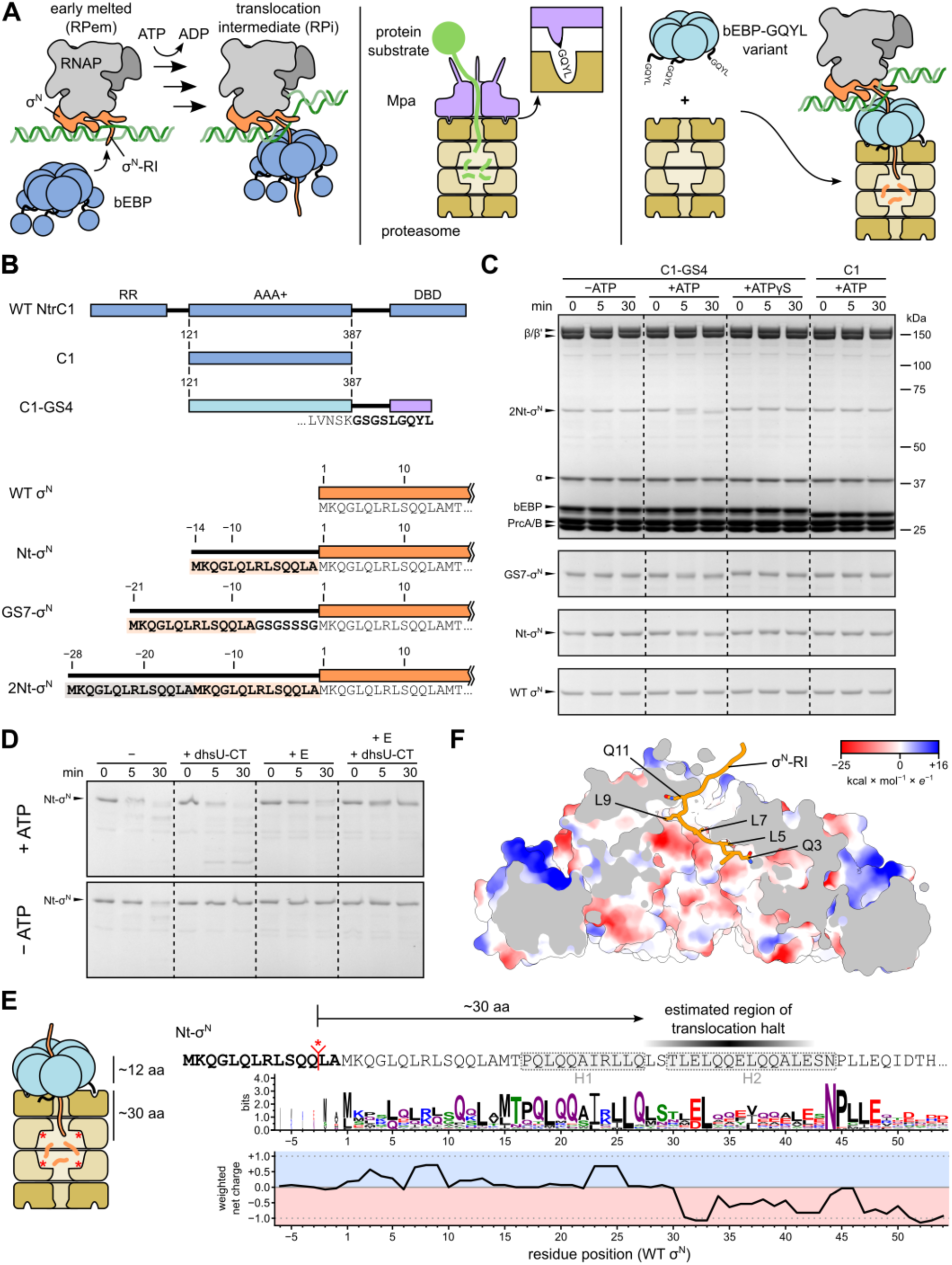
An engineered proteolysis assay measures translocation by the bEBP. (A) The structures suggest that the bEBP (blue; shown with C-terminal domains) actively threads σ^N^-RI (orange) through its pore, extruding σ^N^-RI on the other side (left panel). Compartmentalized proteases such as the mycobacterial proteasome require an AAA+ ATPase (Mpa, purple) to feed protein substrates (green) into the proteolytic chamber (middle panel). Interaction between the proteasome and ATPase is maintained via a short C-terminal motif (GQYL) in the ATPase that inserts into cognate pockets in the proteasome (inset). In our assay design, an engineered variant of the bEBP carrying the GQYL motif (light blue) interacts with the proteasome and feeds σ^N^-RI into the proteolytic chamber. Proteolysis of σ^N^-RI provides a measure for bEBP-mediated translocation during transcription initiation. (B) Schematic representation of the variants used for the proteolytic assay. (C) Distinct band shifts of 2Nt-σ^N^, GS7-σ^N^, and Nt-σ^N^ in presence of ATP and the bEBP variant carrying the GQYL motif (C1-GS4) demonstrate translocation of σ^N^-RI into the proteasome by the bEBP. Reactions contained 0.5 μM RNAP, 0.5 μM σ^N^, 0.6 μM dhsU-CT, and 0.5 μM C1 or C1-GS4 and were carried out at 37 °C. Representative gels of at least two individual experiments are shown. (D) Processive translocation of the bEBP occurs with σ^N^, C1-GS4, and the proteasome - a translocation halt is only observed with the full transcription complex. (E) Estimated lengths required to reach the proteasome active site predict bEBP position on σ^N^-RI during translocation halt (black shading). Position of secondary structure elements of σ^N^-RI present in RPem (H1 and H2) indicated as gray dashed boxes. Experimentally determined cleavage site indicated in red (*) on the sequence of Nt-σ^N^. Net charge distribution of σ^N^-RI changes from positive to negative around residues 28-30. Sequence motif is based on the alignment shown in Extended DAta Fig. 7. See Materials & Methods for details. (F) The surface of the inner pore of the bEBP is lined with negative charges near the hydrophobic pockets that capture residues during translocation. Solvent-excluded surface was clipped to reveal the pore interior and colored according to Coulombic electrostatic potential calculated in ChimeraX.

We appended the five C-terminal residues of Mpa (LGQYL; the proteasome interaction motif) to the C-terminus of C1 with a four-residue Gly-Ser (GS) linker to generate the variant C1-GS4 (Fig. 3B). C1-GS4 showed similar ATPase activity to C1 in the absence or presence of the proteasome (Extended Data Fig. 6A). Compared to C1, transcriptional activation by C1-GS4 was reduced in the absence of the proteasome but strongly increased in the presence of the proteasome (Extended Data Fig. 6B), confirming that C1-GS4 interacts with the proteasome as designed and was functional for activation of Eσ^N^. We further confirmed formation of the complex between C1-GS4 and the proteasome by low-resolution cryo-EM analysis, demonstrating that C1-GS4 caps the proteasome as designed (Extended Data Fig. 6C).

The direct distance of the bEBP pore pocket formed by subunits e-f to the closest active site in the proteasome measures about 90 Å, which corresponds to 26 residues assuming a fully stretched peptide chain with 3.5 Å per residue (i.e., using the N_i_ to N_i+1_ distance in a β-sheet peptide as proxy), or roughly 30 residues accounting for structural constraints in the complex. Therefore, at least 30 residues must be translocated by the bEBP before translocation becomes detectable by proteolysis. To enhance the assay’s readout, we generated variants of σ^N^ with extended N-termini, where 14 N-terminal residues of σ^N^ were duplicated once (Nt-σ^N^), once with a GS linker of 7 residues (GS7-σ^N^; total additional length of 21 residues), or duplicated twice (2Nt-σ^N^; total additional length of 28 residues; Fig. 3B). Each variant was active in a transcription assay but showed decreasing activity with increasing length of the N-terminal extension (Extended Data Figs. 6D and 6E). However, transcription activity of each variant was restored to near wild-type (WT) levels on a promoter template with a pre-melted two-nucleotide bubble (dhsU-CT; Supplementary Table 2; Extended Data Figs. 6D and 6E) previously shown to enhance RPem formation ^33,34^. Thus, the σ^N^ variants with extended N-termini were partially defective in forming RPem but fully functional in subsequent, bEBP-dependent steps of transcription initiation.

Next, we performed the degradation assay with all σ^N^ variants in the context of the full transcription complex – i.e., reactions containing E, WT σ^N^ or σ^N^ variants, dhsU-CT, C1-GS4 and the proteasome – and analyzed the reactions by denaturing gel electrophoresis (Fig. 3C). We did not detect degradation with WT σ^N^ protein, but reactions with Nt-σ^N^, GS7-σ^N^, and 2Nt-σ^N^ showed a small but distinct shift of the σ^N^ band to lower molecular weight. The band shift was observed in the presence of ATP but not with the non-hydrolyzable nucleotide analog ATPγS or C1, confirming that cleavage of σ^N^-RI is dependent on ATP hydrolysis and the ATPase-proteasome interaction. Observation of a distinct band shift, but not a weaker intensity of the band, demonstrates that the bEBP halts translocation of σ^N^ into the proteolytic chamber at a defined position, possibly disengaging from the complex. The translocation/degradation halt was observed only with the full complex (for example, Nt-σ^N^ + E + dhsU-CT promoter fragment); Nt-σ^N^ alone, Nt-σ^N^ + dhsU-CT, or Nt-σ^N^ + E were all completely degraded by the C1-GS4 + proteasome combination in an ATP-dependent manner (Fig. 3D), highlighting the intrinsic processivity of C1.

To determine the cleavage site(s), we subjected the σ^N^ protein bands from reactions in the absence or presence of ATP with WT σ^N^, Nt-σ^N^, and GS7-σ^N^ to N-terminal sequencing by Edman degradation. Reactions without ATP yielded the N-terminal sequence of the full-length proteins (MKQGL; Supplementary Table 3), as expected. Sequencing of the reaction with WT σ^N^ in the presence of ATP also yielded the N-terminal sequence of the full-length protein, suggesting that WT σ^N^ does not reach the active sites of the proteasome during the process of RPo formation. For reactions with Nt-σ^N^, and GS7-σ^N^ in presence of ATP, we identified the N-terminal sequence LA**M**KQGL or LAGSGS, respectively (residue at the WT N-terminus marked in bold). Cleavage in both σ^N^ variants occurred at SQQ/LA, indicating that the GS linker in GS7-σ^N^ was strongly disfavored for proteasomal cleavage, which agrees with the reported substrate preference of the *Mtb* proteasome ^35,36^. We can estimate the point of translocation stop of the bEBP to be roughly 30 residues away from the cleavage site in Nt-σ^N^ (Fig. 3E). Curiously, σ^N^-RI exhibits a conserved switch in local charge environment from slightly positive to negative at this location, and the peptide chain retains an overall negative charge throughout σ^N^-RII (Fig. 3E; Extended Data Fig. 7) ^37^. Furthermore, while the bEBP pore pockets possess a hydrophobic character (Extended Data Fig. 5C), the inner pore surface exhibits a negative electrostatic surface charge (Fig. 3F). During translocation beyond σ^N^-RI E32, the resulting charge repulsion between σ^N^-RI and the environment of DNA and the bEBP inner pore may represent a crucial determinant for bEBP disengagement.

### Pre- and post-catalytic states of bEBP nucleotide hydrolysis suggest a single subunit mechanism

The bEBP engages σ^N^-RI protruding from the DNA bubble with its GAFTGA pore loops, thereby coming in close contact with the DNA. Each bEBP subunit contributes highly conserved residue K213 to form a ring of positive charges around the pore entrance. K213 of alternating bEBP subunits of both RPi1^open^ and RPi1^closed^ (subunits a, c, and e) are positioned to make salt bridge contacts with backbone phosphate oxygens of the DNA, possibly holding the bEBP loosely attached to the DNA during σ^N^-RI translocation. Inside the bEBP pore, hydrophobic pockets formed by residues F216 and T217 of the GAFTGA pore loops and L263 and I203 capture every second residue of σ^N^-RI (Figs. 4A, 4B; Extended Data Figs. 5B and 5C). Notably, σ^N^-RI contains a loosely conserved pattern of residues such that the captured residues are most frequently L or Q (Extended Data Fig. 7).

**Fig. 4:**
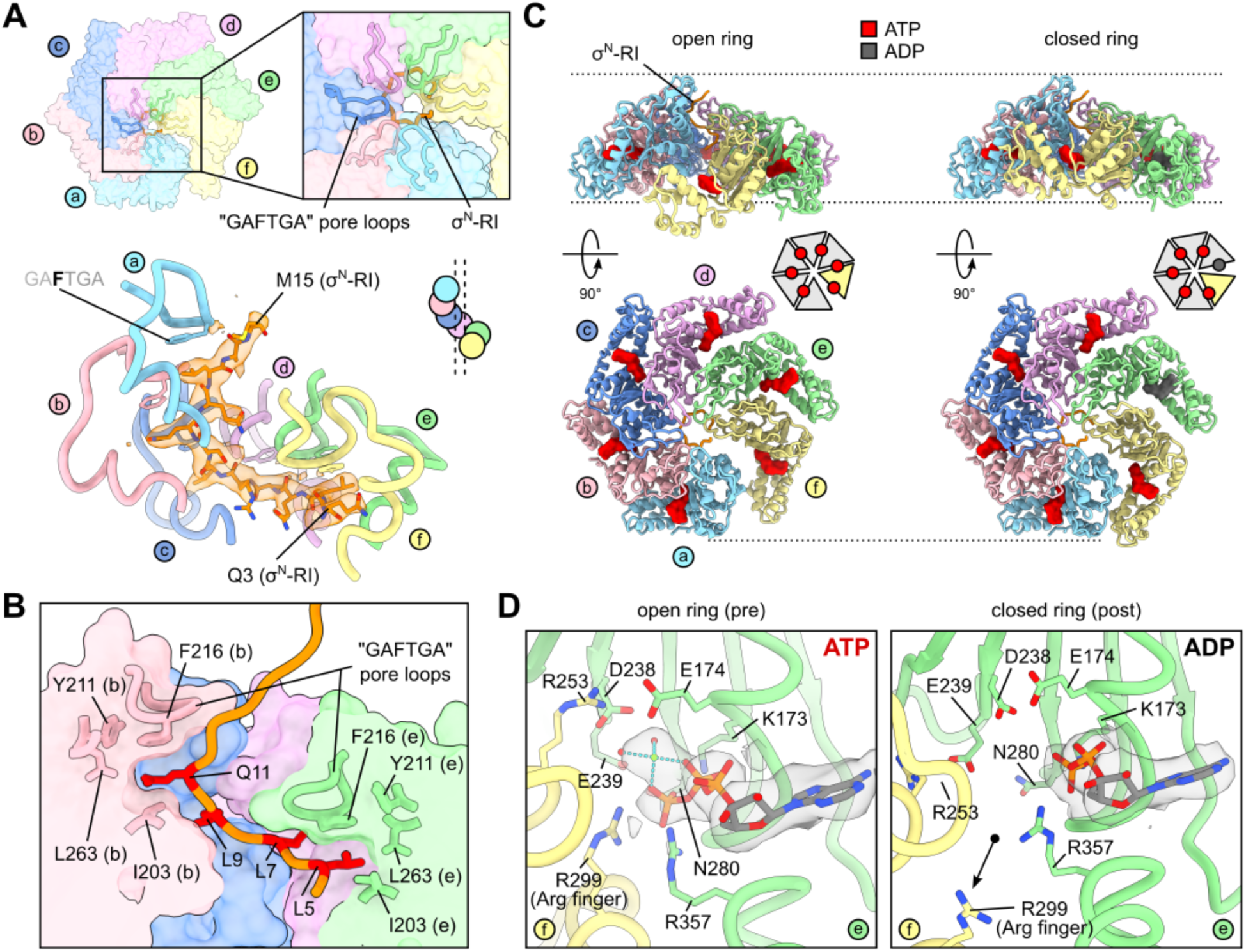
Cryo-EM structures capture intermediate states of bEBP ATP hydrolysis. (A) Conserved “GAFTGA” loops at the bEBP pore (top panel) engage with σ^N^-RI and arrange in a spiral pattern (bottom panel; residues 209-224 of each bEBP subunit and residues 3-15 of σ^N^-RI are shown). Map density for σ^N^-RI (sharpened map) shown as transparent surface (orange). Small inset shows a schematic representation. (B) Clipped surface representation of bEBP subunits reveals captured side chains of σ^N^-RI in hydrophobic pockets. Pocket-forming residues are shown in stick representation. Subunits a and f are hidden for visual clarity. (C) Open and closed ring states correspond to ATP- and ADP-bound states of subunit e. ATP colored in red, ADP colored in grey. The C-terminal domain of subunit e extends below the plane of the hexamer in the ATP-bound state (right), but swings into the plane of the ring in the ADP-bound state (left). Small insets indicate conformational change of subunit f schematically. (D) Active site interactions in subunit e show ATP coordination in a pre-catalytic state (left panel; open ring). Closed ring state contains ADP in the active site of subunit e, corresponding to a post-catalytic state. The arginine finger (R299, f) moved out of the active site (black arrow), and intersubunit salt bridge between E174 (e) and R253 (f) is broken.

In hexameric, ring-forming AAA+ proteins, six ATPase centers are located at the subunit interfaces, where one subunit provides the nucleotide binding pocket while the neighboring subunit contributes essential residues for catalysis ^23^. The RPi1 states are characterized by different conformations of subunit f and different nucleotide occupancies of subunit e (Fig. 4C): In RPi1^open^, a gap of about 10-15 Å exists between subunits a and f and subunit f is rotated out of the ring plane. In RPi1^closed^, the C-terminal domain of subunit f contacts subunit a, thereby bridging the gap, closing the ring and placing subunit f into the plane of the ring. ATP is bound in all active sites of both states, except for the active site of subunit e in RPi1^closed^, which contains ADP (Figs. 4C and 4D; Extended Data Fig. 8). In RPi1^open^, the active site of subunit e is fully formed in a pre-catalytic state poised for catalysis (Fig. 4D): The β- and γ-phosphates of ATP coordinate a Mg^2+^-ion together with active site residues D238 and E239 of subunit e. R299 of subunit f (the “arginine finger”) reaches into the active site contacting the γ-phosphate to provide transition state stabilization during hydrolysis ^38,39^. By contrast, in RPi1^closed^, ADP is bound in the e-f pocket and subunit f is partly dissociated from subunit e, moving the arginine finger out of the active site (Fig. 4D). In both complexes, the active site of subunit a contains an ATP-Mg^2+^ complex situated in a conformation compatible with catalysis. On the other hand, the binding pockets of subunits b, c and d contain ATP in a markedly different conformation (Extended Data Fig. 8): The γ-phosphate of ATP occupies the catalytic Mg^2+^-ion position and is far away from the arginine finger, which is incompatible with the common mechanism of catalysis for AAA+ proteins and C1 specifically ^23,38^. The pre- and post-catalytic structures (RPi1^open^ and RPi1^closed^, respectively) suggest that ATP hydrolysis only occurs in the active site of the second-to-last subunit (e) of the spiral, in agreement with the canonical substrate translocation mechanism of AAA+ proteins ^23^.

### σ^N^-RII occupies the RNAP active site cleft

Density in the RNAP downstream channel of the bEBP-bound classes, including RPi1^open^ and RPi1^closed^, exhibits features consistent with duplex DNA, presumably due to the effective high concentration of DNA ends in the sample, favoring end-binding of DNA. Although this density can be clearly interpreted as duplex DNA, it is comparably weak suggesting heterogeneity within the particle stack. The classification of the bEBP-bound class (Fig. 1C; Extended Data Fig. 2A) using a mask around the bEBP did not reveal any correlation of the ring position with the presence or absence of DNA in the downstream channel. Hence, we performed another 3D classification of the bEBP-bound class focusing on the downstream channel of RNAP (Extended Data Fig. 2B). Approximately half of the classes (corresponding to 43% of the particles) exhibited additional density for σ^N^-RI and σ^N^-RII, and refinement of the best quality classes yielded map “RII” (nominal FSC resolution estimate of 2.7 Å; Extended Data Fig. 9). In the RII complex, residues 52-56 of σ^N^-RI extend over the β’-clamp domain, loosely interacting with the β’ rudder. About 30 residues of σ^N^-RII, including a small helical region (residues 72-75), run along the β’-side of the RNAP cleft and coil between the bridge helix and the gate loop into the vicinity of the active site (Extended Data Fig. 9A). Binding of RII in the downstream channel agrees with the prevalence of negatively charged residues in RII (Extended Data Fig. 7) ^37^. The RII position in these states clashes with the path of the DNA template strand (t-strand) in RPo (Extended Data Fig. 9B), suggesting that RII counteracts nonspecific nucleic acid binding under physiological conditions, analogous to σ^70^ region 1.1 (Extended DAta Fig. 9C) ^40^. RII binding in the downstream channel was also observed in a crystal structure of Eσ^N^ ^41^, and thus appears to remain in the downstream channel through promoter binding and the early steps of initiation.

### σ^N^-RII intertwines with the t-strand DNA in RPo

To gain insight into subsequent steps of RPo formation with σ^N^, we conducted further cryo-EM analysis of the “melted” class (Fig. 1C) by subjecting the particle stack to another 3D classification (Extended Data Fig. 2C). All classes contained a large transcription bubble with density of varying quality for the t-strand. Similar high-quality classes were pooled and refined, yielding two final classes (Figs. 5A and 5B; Extended Data Fig. 2C). The first class (RPo) was refined to a global FSC resolution estimate of 2.8 Å (45,631 particles; Extended Data Fig. 10). In the resulting map, the transcription bubble is opened from position −9 to −1 with the TSS forming the last base pair at the edge of the downstream fork (Figs. 5A and 5C), similar to a previously determined Eσ^N^ RPo structure ^42^. The bases of the small bubble present in RPem (two nucleotides; at positions −11 and −10) and RPi1 (one nucleotide; at position−11) are paired again and contained within the upstream duplex DNA (Fig. 5C). The σ^N^-ELH is wedged into the bubble with conserved W328 located right at the edge of the upstream fork, possibly playing a role in stabilizing the fork. Density for the bEBP and σ^N^-RI are lost entirely, suggesting that the bEBP has disengaged in the RPo state. Strikingly, σ^N^-RII residues 90-120 loop around the DNA t-strand and exit the RNAP main channel between the β flap domain and the β protrusion (Fig. 5D), seemingly acting as an additional measure to stabilize the open transcription bubble and helping to position the t-strand.

**Fig. 5:**
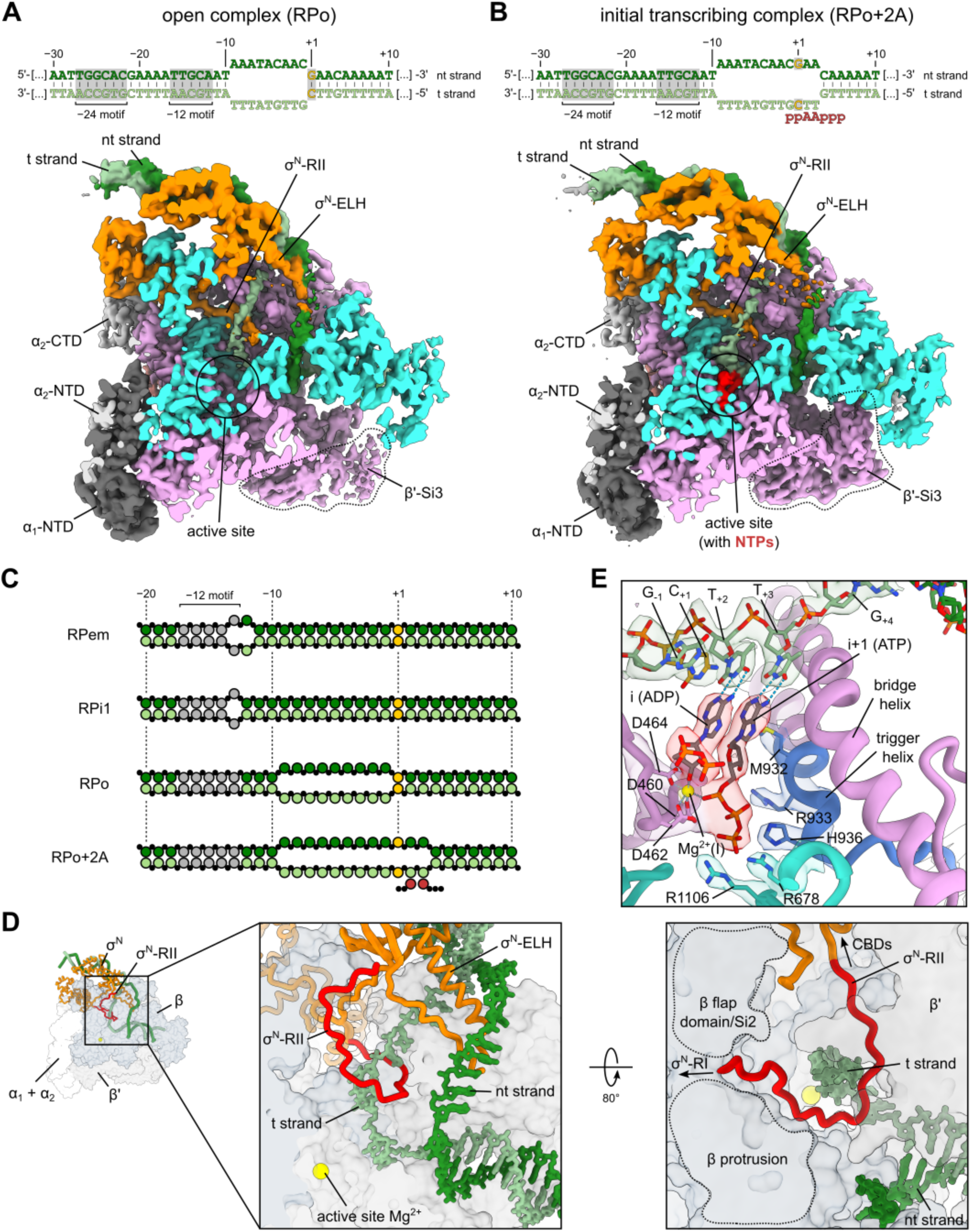
Stabilization of the DNA t-strand by σ^N^-RII in RPo and RPo+2A. (A) Unsharpened map of the RPo complex containing a transcription bubble opened from −9 to −1 (top schematic). (B) Unsharpened map of RPo+2A, an initial transcribing complex containing nucleotides (red) in the active site. The transcription bubble is opened from −9 to +3 (top schematic). (C) Schematic comparison of melted regions in the DNA for complexes along the σ^N^ transcription initiation pathway. (D) In RPo (and RPo+2A), σ^N^-RII (red) loops around the DNA t-strand (pale green; left panel) and exits the complex between β-Si2/flap domain and the β protrusion (right panel). The model of RPo is shown with β/β’ subunits in clipped surface representation to reveal the interior. Labeled arrows indicate neighboring structural elements in σ^N^ for clarification. (E) Active site of RPo+2A contains ADP in the i site and ATP in the i+1 site in a pre-catalytic state. The trigger helix (blue) coordinates triphosphates of ATP (i+1). Density of the sharpened map around DNA and key residues shown as transparent surface.

The 3D classification of the “melted” particle stack yielded an additional, distinct class (RPo+2A; Extended Data Fig. 2C), where density for the typically flexible β’-Si3 domain is well-defined and located in the cleft between β’ and β above the downstream channel (global FSC resolution of 3.1 Å; 15,649 particles; Extended Data Figs. 2C and 11). The resulting map represents an initial transcribing complex (RPitc) in which the transcription bubble is opened from position −9 to +3 (Figs. 5B and 5C). The t-strand is positioned in the RNAP active site and RII loops around the t-strand as seen in RPo, but the position where σ^N^-RII crosses the template strand is shifted by one base (Extended Data Fig. 12). The active site contains two nucleotides – ADP in the “i” site and ATP in the “i+1” site – and the trigger loop is folded, indicating that the complex is poised for catalysis (Fig. 5E). The templating bases positioned in the “i” and “i+1” sites are T+2 and T+3, placing the TSS (C+1; Supplementary Table 1) in the RNAP main channel. The conditions of complex preparation for cryo-EM included a vast excess of ATP (5 mM) as the sole source of nucleotide, presumably favoring the observed start site selection.

## Discussion

Through functional and structural analysis, we show that the bEBP exerts limited ATP-powered mechanical action on σ^N^-RI to pull it through the initial DNA bubble of RPem, thereby disrupting the σ^N^-RI:ELH inhibitory interactions. Translocation of nucleic acids or proteins by AAA+ remodellers occurs typically with high processivity. Although the bEBP displays processivity in the absence of the full transcription complex (degrading the entire 477 amino acid length of σ^N^; Fig. 3D), in the presence of the full complex translocation of σ^N^ occurs only for about 15 rounds of ATP hydrolysis, corresponding to roughly 30 residues (Fig. 3E). The local negative net charge in σ^N^-RI in this region may create sufficient repulsion between the DNA and/or the inner pore surface of the bEBP to cause the translocation stop. To disengage, the bEBP ring may fall off as a complex or dissemble into smaller oligomers or even individual subunits. Therefore, Eσ^N^ transcription activation represents the bacterial counterpart of an exclusive group of examples found in eukaryotes, where the mechanistic principle of activation by limited substrate translocation by an AAA+ ATPase is utilized: Mitochondrial ClpX activates the initiating enzyme for heme biosynthesis by partial unfolding to enable cofactor insertion ^43^; TRIP13 remodels the cell-cycle protein MAD2 of the mitotic checkpoint complex by local unwinding at its N-terminus to trigger completion of the spindle assembly checkpoint ^44,45^; Rubisco activase rescues the photosynthetic enzyme complex from sugar phosphate inactivation by selective engagement and local destabilization at the C-termini of the large subunits ^46^; NSF recycles membrane fusion SNARE proteins in a single step of ATP hydrolysis, representing the lower extreme of limited translocation ^47,48^.

Determining the kinetics of RPo formation (Figs. 1A and 1B) enabled timing of the cryo-EM sample preparation to obtain structures of bEBP-bound Eσ^N^ initiating complexes in the presence of ATP, including intermediates of RPo formation (Figs. 1C and 2). The structures caught the bEBP in action while mechanically unfolding σ^N^-RI, demonstrating that the bEBP works as a protein remodeller, consistent with the function of many AAA+ proteins ^23^. Furthermore, we engineered a proteolysis assay which was used to demonstrate biochemically that the bEBP exhibits substrate translocation and unfolding activity (Fig. 3), firmly supporting the structural data. Within the bEBP-bound structures, the majority of particles were intermediate complexes where the bEBP advanced by one step (i.e., translocation of σ^N^-RI by two residues; RPi1), which was sufficient to partially unfold H1 of σ^N^-RI. The interaction between σ^N^-RI and the ELH remained intact in these intermediates (Fig. 2B). No further advanced intermediates could be determined from our dataset, indicating that disrupting the σ^N^-RI:ELH interaction represents a strong energetic barrier and a rate-limiting step for RPo formation. The lack of intermediate states between RPo and RPi1 in our data can be explained by a highly stabilized RPo state corresponding to a large free energy change ^28^ resulting in sparsely populated intermediates following the rate-limiting step.

Determining which and how many subunits perform nucleotide hydrolysis at each step of substrate translocation in AAA+ proteins is crucial to understanding their mechanism. Our structures suggest that the bEBP works according to the canonical hand-over-hand substrate translocation mechanism of AAA+ proteins ^23^ as follows (Fig. 6A): At each step, only subunit e is catalytically active and hydrolyses ATP. Upon ATP cleavage, the last subunit (f) can dissociate, bind subunit a, and take the place of subunit a at the top of the spiral. The GAFTGA loop of subunit f would reach up to the protein substrate (σ^N^-RI) and engage with the next residue of σ^N^-RI, thereby advancing the bEBP along the substrate by one step. Highly conserved residue K213 (located right before the GAFTGA motif, i.e., …**K**GAFTGA…) may aid in repositioning the loop for substrate capture by interacting with the DNA phosphate backbone. Subunit e takes the position of the last subunit (f), ADP is exchanged for ATP, and the complex is ready to enter the next cycle of ATP hydrolysis and substrate translocation. Notably, in the post-catalytic state (RPi1^closed^), the C-terminal domain of subunit f has already established contact with subunit a, but the GAFTGA loop has yet to move to the top of the spiral. Likely, we captured this state because this intermediate has piled up at the energetic barrier of the σ^N^-RI:ELH interaction.

**Fig. 6:**
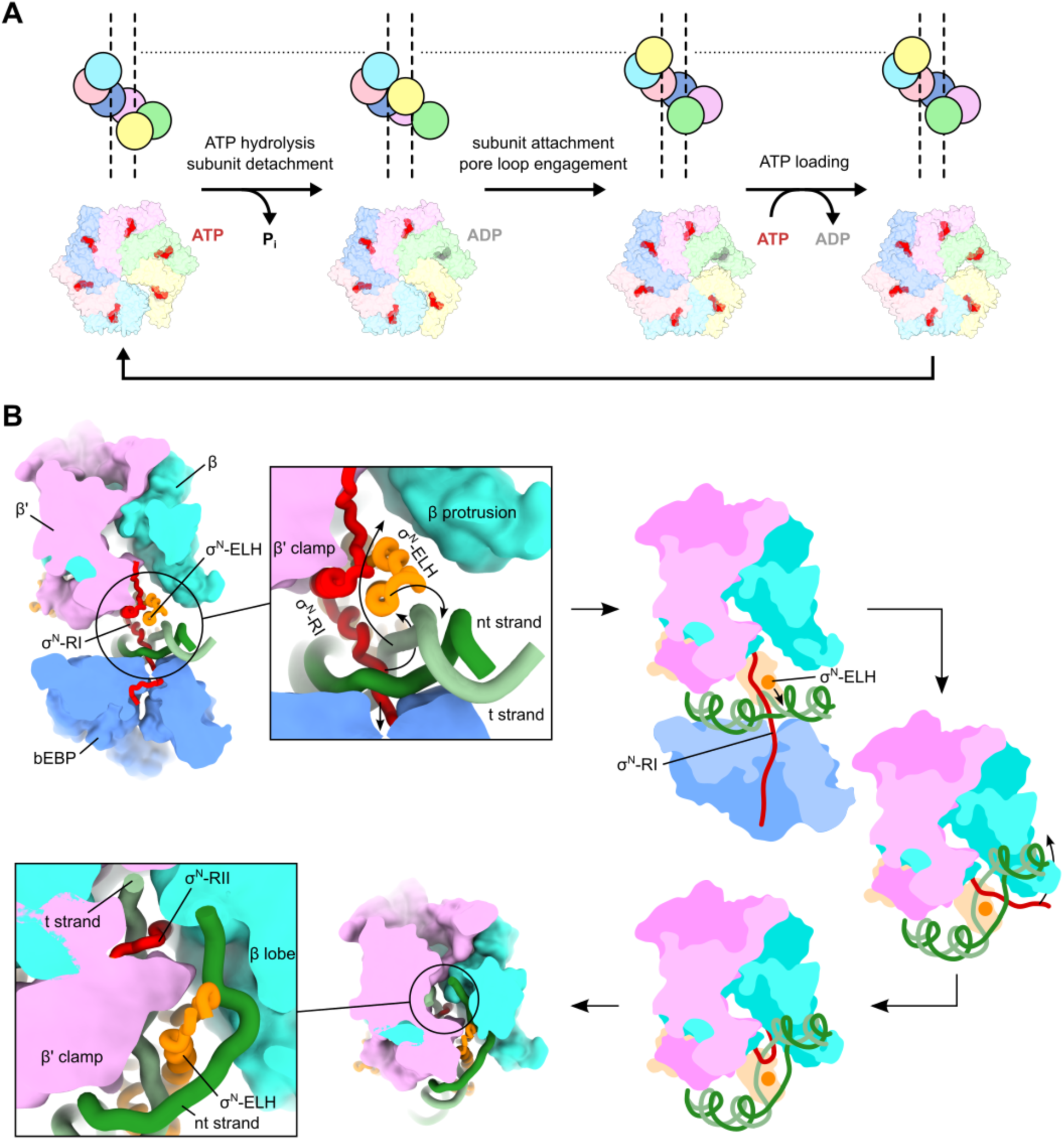
Schematic of the proposed mechanism. (A) For ATP hydrolysis-coupled substrate translocation by the bEBP, the ATP-containing bEBP binds to an unstructured end of the protein substrate (σ^N^-RI) to engage in a spiral conformation, leaving a gap between the top (blue) and bottom (yellow) subunit. ATP hydrolysis in the green subunit disrupts the intersubunit interaction between green and yellow allowing the yellow subunit to reach across the gap and contact the blue/top subunit. The pore loop of the yellow subunit engages with the substrate, and thereby, the yellow subunit assumes the position at the top of the spiral and the complex advanced one step along the substrate. Subsequently, ADP in the green subunit is exchanged for ATP and cycle can start over. (B) For the isomerization from RPi1 to RPo, the transcription initiation complex is viewed along the DNA axis and large subunits (β, β’, bEBP) are clipped along the axis of σ^N^-RI (red) for visual clarity. During translocation of σ^N^-RI by the bEBP, the ELH must insert at the minor groove near the DNA bubble to place the DNA t-strand between σ^N^-RI and the ELH. To loop around the t-strand, σ^N^-RI/RII passes around the ELH and inserts between β-Si2 and the β protrusion forming the arrangement observed in the RPo and RP+2A structures.

A model for the steps that must occur during the transition from RPi1 to RPo can be deduced (Fig. 6B): As the bEBP continues translocation of σ^N^-RI, disrupting the σ^N^-RI:ELH interaction, the ELH must insert between the DNA strands at the minor groove around position −8 to bring the t-strand between σ^N^-RI and the ELH. Subsequently, σ^N^-RI/RII must pass in front of the ELH in order to insert between the β-Si2/flap domain and the β protrusion to arrive at the looped arrangement observed in RPo. Experiments with slowly hydrolysable ATPγS suggest that the bEBP is required for additional steps after disruption of the σ^N^-RI:ELH interaction ^49^, which may be achieved early in the translocation process. It seems plausible that continued bEBP threading of σ^N^ until the translocation halt is required to pull RII out of the RNAP main channel. Thereafter, the RNAP downstream channel is free to engage with DNA, and binding free energy may drive the insertion of the downstream duplex DNA into the channel.

A remarkable feature of the RPo and RPo+2A/RPitc structures is the looping of σ^N^-RII around the t-strand DNA (Fig. 5D). In Eσ^70^ transcription complexes, σ^70^ region 3.2 (the “σ-finger”) – which occupies a similar location in RPo as σ^N^-RII ^50,51^ – facilitates efficient formation of the first phosphodiester bond ^52,53^, presumably by helping to position the t-strand DNA. The RII:t-strand junction may fulfill a similar role and help to position the t-strand for initial catalysis. In recently determined RPitc structures with different lengths of nascent RNA the RII loop lies beneath the t-strand and is not topologically entangled ^54^. However, these samples were prepared for structural analysis by assembling complexes with promoter DNA containing a pre-melted bubble, a “bypass” variant of σ^N^, and in the absence of a bEBP and so do not reflect *bona fide*, bEBP-dependent Eσ^N^ RPitc complexes.

The entanglement of RII with the t-strand is a topological conundrum that must be resolved before RNAP can leave the promoter. Regulated bEBP disassembly prevents the bEBP from having to unfold the entire length of the σ^N^ polypeptide (477 residues) through the DNA bubble and presumably allows σ^N^ to disentangle from the DNA t-strand (Fig. 5D) as RNAP escapes the promoter. Further work will be required to elucidate the structural details for the steps of RPo formation following RPi1, and for how the entanglement of σ^N^ with the DNA bubble is resolved for promoter escape.

## Acknowledgments and funding sources

We are grateful to B. Tracy Nixon, Ruth Saecker, Brandon Malone, Ruby Froom, James Chen, Elizabeth Campbell, and Robert Landick for constructive discussion and critical reading of the manuscript. We thank Eilika Weber-Ban (ETH Zurich, Switzerland) for the generous gift of the expression plasmid of the *M. tuberculosis* open-gate proteasome. We thank Johanna Sotiris, Honkit Ng, and Mark Ebrahim of the Evelyn Gruss Lipper Cryo-Electron Microscopy Resource Center (The Rockefeller University, New York, NY 10065, USA) for support with electron microscopy instrumentation and sample handling. We thank Michael Berne of the Tufts University Core Facility (Tufts Medical School, Boston, MA 02111, USA) for the Edman degradation-based protein sequencing service. A.U.M. is an Agouron Institute Awardee of the Life Sciences Research Foundation. This work was supported by a grant from the NIH to S.A.D. (R35 GM118130).

## Author contributions

A.U.M. and S.A.D. conceived and designed the study. A.U.M. and N.M. prepared purified proteins and performed biochemical experiments. A.U.M. performed cryo-EM sample preparation, data collection, processed and analyzed the data. A.U.M. and S.A.D. interpreted structural data and prepared models. A.U.M. drafted the manuscript and prepared figures. A.U.M. and S.A.D. wrote the full manuscript. All authors provided comments and contributed to editing.

## Materials and Methods

### Protein expression and purification

All purification steps were performed at 4°C or on ice if not stated otherwise.

#### Eco core RNAP

Full-length *Eco* RNAP (UniProt entries P0A7Z4, P0A8V2, P0A8T7, P0A800 for α, β, β’, and ω) was expressed and purified largely following the procedure as described previously ^55^. Specifically, a pET-based plasmid harboring full length RNAP subunits α, β, ω as well as β’-PPX-His_10_ (PPX; PreScission protease site, LEVLFQGP, Cytiva) under an IPTG-inducible promoter (addgene #128940) was co-transformed with pACYC-Duet-1 encoding ω (addgene #128837) into *Eco* BL21(DE3). Shaking cultures of a total of 6 L LB media supplemented with 100 μg/ml ampicillin and 34 μg/ml chloramphenicol were inoculated with an overnight preculture from freshly transformed cells and grown to an OD600 of 1 at 37°C. Expression was induced with IPTG (0.5 mM final concentration) and cultures were incubated for 3 hr at 30°C. Cells were harvested by centrifugation (Beckman JS-4.2, 4,500xg, 4°C, 20 min) and pellets were resuspended in a total of 120 mL lysis buffer (50 mM Tris-HCl, pH 8/RT, 10 mM DTT, 1 mM ZnCl_2_, 5% (v/v) glycerol, 0.5x c0mplete EDTA-free protease inhibitors (Roche), 1 mM PMSF). Resuspended cells were stored at −80°C until use. Cell suspensions were thawed, lysed by high-pressure shearing (Avestin EmulsiFlex C50), and insoluble material was removed by centrifugation (Beckman JA-20, 27,000xg, 4°C, 30 min). While stirring, polyethyleneimine [PEI; 10% (w/v) in water adjusted to pH 8/RT with HCl] was slowly added to a final concentration of 0.6% (w/v) to the soluble fraction and the mixture was incubated for 25 min at 4°C while stirring. After collection of the PEI precipitate by centrifugation (Beckman JA-20, 27,000xg, 4°C, 1 hr), the pellets were washed three times in a total of 210 mL PEI wash buffer [50 mM Tris-HCl, pH 7.9/RT, 0.5 M NaCl, 5% (v/v) glycerol, 10 mM DTT]. For each wash step, the pellets were resuspended using a glass Dounce homogenizer (Wheaton) and the PEI precipitate was collected again by centrifugation. To elute RNAP from the PEI, the pellets from the last wash step were resuspended as above in a total of 120 mL PEI elution buffer [50 mM Tris-HCl, pH7.9/RT, 1 M NaCl, 5% (v/v) glycerol, 10 mM DTT] and the PEI precipitate was collected again by centrifugation saving the supernatant. Elution was repeated for a total of three rounds pooling the supernatant from each centrifugation step (final total of 360 mL). 35 g/100 mL ammonium sulfate (grounded to powder) were added slowly while stirring and the mixture was incubated overnight at 4°C. Ammonium sulfate precipitate was recovered by centrifugation (Beckman JA-18, 33000xg, 4°C, 30 min) and resuspended in a total of 50 mL IMAC buffer A [20 mM Tris-HCl, pH 7.8/RT, 1 M NaCl, 5% (v/v) glycerol, 1 mM β-mercaptoethanol]. The sample was passed through 5 μm filter and applied to 2×5 mL HiTrap IMAC HP columns (Cytiva) connected in series, charged with Ni^2+^ and equilibrated in IMAC buffer A. After washing the columns with increasing imidazole concentrations in buffer A (last wash with 80 mM imidazole), protein was eluted in buffer B [20 mM Tris-HCl, pH 7.8/RT, 1 M NaCl, 250 mM imidazole, 5% (v/v) glycerol, 1 mM β-mercaptoethanol]. Fractions were pooled according to protein content and His-tagged 3C protease was added at 1:36 molar ratio. The sample was dialyzed overnight at 4°C in a 12-14 kDa cutoff membrane (SpectraPor) against 20 mM Tris-HCl, pH8/4°C, 1 M NaCl, 5% (v/v) glycerol, 0.1 mM EDTA, 1 mM β-mercaptoethanol, 0.5 mM DTT. After dialysis, the sample was passed over the IMAC columns followed by dialysis of the flow-through overnight at 4°C in a 12-14 kDa cutoff membrane (SpectraPor) against 10 mM Tris-HCl, pH7.8/RT, 100 mM NaCl, 0.1 mM EDTA, 5% (v/v) glycerol, 5 mM DTT. After dialysis, the sample was loaded on a 40 mL Biorex column (Biorad Biorex-70 resin #142-5842) equilibrated in Biorex A buffer [10 mM Tris-HCl, pH7.8/RT, 0.1 mM EDTA, 5% (v/v) glycerol, 5 mM DTT] and eluted with a gradient of 20-80% Biorex B buffer [10 mM Tris-HCl, pH 7.8/RT, 1 M NaCl, 0.1 mM EDTA, 5% (v/v) glycerol, 5 mM DTT] over 15 column volumes. Fractions were pooled according to protein content. The pool was concentrated using a centrifugal filter (3,500xg, 4°C, Amicon Ultra 100K, EMD Millipore) and loaded on a Superdex 200 26/600 320 mL column (Cytiva) equilibrated in 20 mM HEPES-Na, pH 7.8/RT, 0.5 M NaCl, 0.1 mM EDTA, 5% (v/v) glycerol, 0.5 mM TCEP. Peak fractions were pooled, concentrated using centrifugal filters (3,500xg, 4°C, Amicon Ultra 100K, EMD Millipore) and mixed with buffer containing 50% (v/v) glycerol to achieve a final concentration of 20% (v/v) glycerol. Aliquots were frozen in liquid N_2_ and stored at −80°C until use.

#### Eco σ^N^ and variants

Tagless, full-length *Eco* σ^N^ (RpoN, Sig54; UniProt entry P24255) was purified as described previously ^21^. Variants of σ^N^ were obtained by cloning full-length σ^N^ into pET28a with an N-terminal His_10_-SUMO tag and introducing the desired mutations by PCR amplification with oligonucleotide primers carrying the desired modifications and ligation of the PCR product. Constructs were freshly transformed into *Eco* BL21(DE3) cells and transformants were grown at 37°C in 200 mL LB shaking cultures (baffled flasks) containing 50 ug/ml kanamycin. At OD600 of 0.6, the cultures were equilibrated to 16°C for 30 min before induction with 1mM IPTG. Expression was carried out overnight at 16°C. Cells were harvested by centrifugation (Beckman JS-4.2, 4,500 xg, 4°C, 30 min) and resuspended in lysis buffer [20 mM Tris-HCl, pH8/RT, 500 mM NaCl, 5 mM imidazole, 5% (v/v) glycerol, 0.5 mM DTT, 1 mM PMSF, 1x c0mplete EDTA-free protease inhibitor (Roche)]. At this point, the cell resuspensions were frozen in liquid N_2_ and stored at −80°C until further processing. After thawing, cells were lysed by sonication (Branson Digital Sonifier 450) and subjected to centrifugation (Beckman JA-20, 11,000 xg, 4°C, 1 hr) to remove insoluble material. The soluble fraction was passed through 0.45 μm filter and applied to a 1 mL Ni^2+^-charged IMAC HiTrap (Cytiva) equilibrated in buffer A [20 mM Tris-HCl, pH8/RT, 500 mM NaCl, 5% (v/v) glycerol, 0.5 mM DTT]. The column was washed with increasing concentrations of imidazole in buffer A (40 mM, 80 mM, 120 mM, 350 mM imidazole) while collecting fractions. Fractions containing *Eco* σ^N^ protein were pooled; His-tagged Ulp1 protease was added at 1:40 molar ratio and the sample was dialyzed overnight at 4°C against buffer A in a 12-14 kDa cutoff membrane (SpectraPor). After recovering the sample from dialysis, it was passed again over the IMAC resin to remove uncleaved fusion protein and the Ulp1 protease. The flow-through was collected, concentrated using centrifugal filters (3,500xg, 4°C, Amicon Ultra 30K, EMD Millipore) and mixed with an equal volume of storage buffer [20 mM Tris-HCl, pH8/RT, 500 mM NaCl, 40% (v/v) glycerol, 0.1 mM EDTA, 2 mM DTT]. Aliquots were frozen in liquid N_2_ and stored at −80°C until use.

#### Aae C1 (bEBP) and variants

The constitutively active variant of *Aae* NtrC1 (UniProt entry O67198) missing the N- and C-terminal domains (C1; residues 121-387 of the full-length protein) was purified as described previously ^21^. An expression construct for C1 carrying the proteasome interaction motif (C1-GS4) was obtained by amplification of the parent expression vector using mutagenic primers (Supplementary Table 2) to encode residues GSGSLGQYL at the C-terminus and ligation of the linearized product. Purification of C1-GS4 was carried out in analogy to the procedure described for C1 but required several modifications: Specifically, freshly transformed *Eco* BL21(DE3) were grown in 2 L LB medium supplemented with 100 μg/ml ampicillin as a shaking culture at 37°C. Expression was induced with 1 mM IPTG at an OD600 of 0.6 and continued overnight at 25°C. Cells were harvested by centrifugation (Beckman JS-4.2, 4,500xg, 4°C, 20 min) and pellets were resuspended in lysis buffer [50 mM Tris-HCl, pH 8/RT, 500 mM KCl, 5% (v/v) glycerol, 5 mM EDTA, 1 mM TCEP] supplemented with 1x c0mplete EDTA-free (Roche) and 1 mM PMSF. After cell lysis by high-pressure shearing (Avestin EmulsiFlex C50), insoluble material was removed by centrifugation (Beckman JA-18, 30,000xg, 4°C, 30 min). The supernatant was recovered and placed in a water bath equilibrated at 70°C for 30 min while stirring. Following removal of the precipitate by centrifugation (Beckman JA-18, 30000xg, 4°C, 30 min), protein was precipitated with ammonium sulfate at 60% saturation at 4°C for 1 hr. Precipitate was collected by centrifugation (Beckman JA-18, 30,000xg, 4°C, 30 min) and resuspended in 20 mM Tris-HCl, pH 8/RT, 500 mM KCl, 1 mM EDTA, 1 mM DTT. The resuspension was dialyzed overnight at 4°C in a 12-14 kDa cutoff membrane (SpectraPor) against 20 mM Tris-HCl, pH 8/RT, 10 mM KCl, 1 mM EDTA, 1 mM DTT. Subsequently, the sample was loaded on a 5 mL HiTrap Q HP column (Cytiva) equilibrated in low salt buffer (20 mM Tris-HCl, pH 8/RT, 50 mM KCl, 1 mM EDTA, 1 mM DTT) and eluted with a salt gradient of 50 mM to 500 mM KCl over 15 column volumes. Fractions were pooled according to protein content and dialyzed overnight at 4°C in a 12-14 kDa cutoff membrane (SpectraPor) against 20 mM MES, pH 6.5/RT, 10 mM KCl, 1 mM DTT. The dialyzed sample was loaded on a 5 mL HiTrap SP HP column (Cytiva) and eluted with a salt gradient of 50 mM to 300 mM KCl over 20 column volumes. Fractions were pooled according to protein content and concentrated using centrifugal filters (3,500xg, 4°C, Amicon Ultra 30K, EMD Millipore). Aliquots were frozen in liquid N_2_ and stored at −80°C until use.

#### Bacterial proteasome

The “open-gate” variant of the 20S core particle of the *M. tuberculosis* proteasome (20S-og) was expressed recombinantly in *Eco* BL21(DE3) from a pETDuet-1 vector encoding prcAΔN7 and prcB with a C-terminal Strep tag (WSHPQFEK). The expression construct was a gift from Eilika Weber-Ban (ETH Zurich, Switzerland). Cells were grown in ZYP-5052 autoinduction media as three 2 L shaking cultures at 25°C overnight and harvested by centrifugation (Beckman JS-4.2, 4,500xg, 4°C). Pellets were resuspended in buffer P [50 mM Tris-HCl, pH7.8/RT, 150 mM NaCl, 1 mM EDTA, 10% (v/v) glycerol, 1 mM DTT) supplemented with 1 mM DTT (final 2 mM), and cells were lysed by high-pressure shearing (Avestin EmulsiFlex C50). Insoluble material was removed by centrifugation (Beckman JA-18, 33,000xg, 4°C, 1 hr). The supernatant was recovered, passed through a 5 μm filter, and applied to a 5 mL StrepTactin XT 4Flow high-capacity resin (IBA Lifesciences) equilibrated in buffer P. After washing the resin with 5 column volumes of buffer P, the protein was eluted with buffer A containing 50 mM biotin. Elution fractions were pooled according to protein content and further purified by gel filtration (Superdex 200 HiLoad 26/600, 320 mL, Cytiva) in buffer P. Peak fractions were pooled, concentrated to 15-30 μM proteasome complex using centrifugal filters (3,500xg, 4°C, Amicon Ultra 100K, EMD Millipore) and stored at 4°C. Aliquots of the sample were applied to a Superose 6 10/300GL (24 mL) column (Cytiva) in buffer P before use to remove large aggregates. Peak fractions were pooled, concentrated and stored at 4°C.

#### TSS determination by 5’-RACE with template switching

We performed rapid amplification of cDNA ends (RACE) including reverse transcription with template switching ^56^ to determine the 5’-end sequences of *in vitro* transcribed RNAs from the linear dhsU and dhsU+2T promoter fragments (−60 to +30; Supplementary Table 2). Promoter DNA fragments were annealed from synthetic oligos to form dhsU (oligos dhsU_top and dhsU_bot; Supplementary Table 2) and dhsU+2T (oligos dhsU+2T_top and dhsU+2T_bot; Supplementary Table 2) duplex DNA. Oligos were mixed at 2 μM final concentration in 10 mM HEPES-NaOH, pH 8/RT, 50 mM NaCl and subjected to the following program in a thermocycler: 95°C for 30 s, 95°C for 15 s for 70 cycles with −1°C/cycle, 25°C hold. Reactions containing 80 nM RNAP, 200 nM σ^N^, 20 nM promoter DNA, 1 μM C1, 1 mM of each CTP/GTP/UTP, 5 mM ATP were assembled in a 120 μL reaction in buffer TXN (40 mM Tris-HCl, pH 8/RT, 200 mM KCl, 10 mM MgCl_2_, 5 mM DTT, 40 μg/ul BSA) as follows: RNAP and σ^N^ were mixed and incubated for 10 min at 37°C to form holoenzyme; DNA was added and the reaction was incubated for 15 min at 37°C to form the early-melted intermediate (RPem); C1 was added and the reaction was incubated for 5 min at 37°C before starting the reaction with the addition of NTPs. After incubating the reaction for 30 min at 37°C, 2 μL TURBO DNase (ThermoFisher Scientific #AM1907; final concentration 33 U/mL) was added and the reaction was incubated for 15 min at 37°C to degrade the promoter DNA. To eliminate free Mg^2+^ ions, 5 μl of 0.5 M EDTA were added (final concentration 19.7 mM) and the reaction was placed on ice. RNA was purified using the Oligo Clean & Concentrator kit (Zymo #D4060), eluting in 20 μL of 20 mM Tris-HCl, pH 8/RT. RNA preparations were subjected to capping by the Vaccinia virus Capping Enzyme (VCE; NEB #M2080S) in 20 μL reactions containing 0.5 mM GTP, 1 U/μL RNase inhibitor (NEB #M0314S), 0.5 U/ul VCE in 1x capping buffer (NEB), and 12 μL purified RNA. RNA was heat-denatured at 70°C for 5 min and cooled on ice before addition to the reaction. Capping reactions were incubated for 2 hr at 37°C and treated RNAs were purified using the Oligo Clean & Concentrator kit (Zymo #D4060), eluting in 20 μL water. 4 μL purified capped RNAs were mixed with RT primer (dhsU-RTprim; Supplementary Table 2; final concentration 1 μM) and dNTP mix (NEB #N0447S; final concentration 1 mM) in 6 μL total volume, denatured at 70°C for 5 min and placed immediately on ice. A working solution of reverse transcriptase and template switching (RTTS) reagents (NEB #M0466S) was prepared by mixing 2.5 parts template switching RT buffer, 1 part of 10 μM RACE-TSO_fw primer (Supplementary Table 2), and 1 part of RT enzyme mix. 4 μL of RTTS mix were added to the denatured RNA-RTprimer-dNTP mix (total volume 10 μL). The reaction was incubated in a thermocycler at 42°C for 90 min, heated to 85°C for 5 min, and cooled to 4°C. Reactions were diluted two-fold with water, and 2.5 μL of diluted RTTS reaction were used in 25 μL cDNA amplification reactions. PCR mixes contained 0.2 mM dNTPs, 0.5 μM RACE-PCR-TSO_fw primer (Supplementary Table 2), 0.5 μM RACE-PCR-RT_rv primer (Supplementary Table 2), 0.02 U/μL Q5 HotStart polymerase (NEB #M0493S) in Q5 reaction buffer (NEB) and were subjected to the following PCR program: initial denaturation at 98°C for 30 s; 5 cycles of 98°C for 10 s, 72°C for 2 s; 5 cycles of 98°C for 10 s, 70°C for 2 s; 35 cycles of 98°C for 10 s, 65°C for 15 s, 72°C for 15 s; final extension at 72°C for 30 s; hold at 10°C. Two 25 μL cDNA amplification reactions were performed for each promoter DNA. Free primers were digested by adding 0.5 μL (10 U) of Exonuclease I (NEB #M0293S) to each reaction and incubating the reactions at 37°C for 30 min. PCR products were purified using the Oligo Clean & Concentrator kit (Zymo #D4060), eluting in 12 μL 20 mM Tris-HCl, pH 8/RT. Purified PCR products were cloned into a pTwistAmp vector (Twist Bioscience) linearized using primers RACE-pTwist_fw and RACE-pTwist_rv (Supplementary Table 2) by isothermal DNA assembly (NEB #E5520S). Assembly was performed by mixing 15 fmol linearized vector with 90 fmol cDNA PCR product in 2.5 ul total volume. An equal volume of 2x assembly enzyme mix (NEB #E5520S) was added and reactions were incubated at 45°C for 30 min. 3 μL of the assembly reactions were transformed into 50 μL chemo-competent NEB5a cells (NEB #E5520S) according to the manufacturer’s protocol. Recovered transformants were plated on LB agar plates containing 100 μg/ml ampicillin, and plates were incubated at 37°C overnight. Plates were subjected to direct colony sequencing (Azenta/Genewiz, New Jersey, USA) of 10 random clones for each promoter DNA fragment using the M13FOR primer (Supplementary Table 2). Sequencing results were aligned using ClustalOmega and transcription start sites were identified as being directly preceded by 4 G bases (for capped RNAs) or 3 G bases (for uncapped RNAs) at the 3’-end of the RACE-TSO_fw sequence as previously described ^56^. Nine out of nine sequencing reactions yielded GAACAA as the 5’-end sequence for dhsU (Supplementary Table 1). Six out of seven sequencing reactions yielded GTACAA as the 5’-end sequence for dhsU+2T. Failed sequencing reactions and transcription via end binding were eliminated from the analysis. The results established the +2 site, and that introduction of a T at this position did not change the TSS.

#### Stopped flow fluorescence spectroscopy for RPo formation kinetics

A dhsU promoter fragment labeled with Cy3 at the +2 position (Cy3-dhsU+2T) was prepared by annealing 10 μM dhsU+2T-Cy3_top (Supplementary Table 2) and 11 μM dhsU+2T_bot (Supplementary Table 2) in 10 mM HEPES-Na, pH 8/RT, 50 mM NaCl in a thermocycler (program: 95°C for 30 s, 95°C for 15 s for 70 cycles with −1°C/cycle, 25°C hold). Reaction mix A was prepared in buffer S (40 mM Tris-HCl, pH 8/RT, 200 mM KCl, 10 mM MgCl_2_, 1 mM DTT, 10 μg/ml BSA) by first mixing RNAP and σ^N^ and incubating for 5 min at 37°C; Cy3-dhsU+2T was added and the reaction was incubated for 10 min at 37°C; C1 was added and the reaction was incubated for 2 min at 37°C. Reaction mix B was prepared by mixing ATP in buffer S. Reaction mixes A and B were loaded into a stopped flow instrument (Applied Photophysics SX20; 535 nm LED, 550 nm shortpass excitation filter, 570 longpass emission filter) equilibrated at 37°C and incubated 1 min before starting the experiment. At least 5 traces were recorded and averaged. For analysis, the averaged traces were offset corrected by subtracting the value of the first data point from all data points within a series. Final concentrations after mixing in the stopped flow cell were 50 nM RNAP, 75 nM σ^N^, 10 nM Cy3-dhsU+2T, 250 nM C1, 2.5 mM ATP for traces shown in Fig. 1A and Extended Data Fig. 1A; 100 nM RNAP, 150 nM σ^N^, 20 nM Cy3-dhsU+2T, 500 nM C1, 5 mM ATP for traces shown in Extended Data Fig. 1B; 200 nM RNAP, 300 nM σ^N^, 20 nM Cy3-dhsU+2T, 500 nM C1, 5 mM ATP for traces shown in Extended Data Fig. 1C. Fluorescence signal F in the experiment was fit with a single exponential equation (i.e., unimolecular conversion) as follows

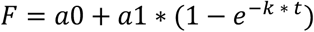

where a0 is the background signal, a1 is the maximum signal, and k corresponds to the apparent rate constant. Parameter estimates from the data fit are presented in Supplementary Table 4. Data fitting was performed with the GraphPad Prism v9.5.1 software.

### Preparation of Eα^N^dhsU-bEBP *de novo* complexes for cryo-EM analysis

#### dhsU promoter DNA

Commercially synthesized DNA oligos (Integrated DNA Technologies) dhsU_top and dhsU_bot (Supplementary Table 2) were resuspended in nuclease-free H_2_O. Equimolar amounts of each strand were mixed and annealed in 10 mM HEPES-NaOH, pH 8.0, 50 mM NaCl at a final duplex concentration of 120 μM.

#### Sample preparation

Purified proteins were desalted into reaction buffer (40 mM Tris-HCl, pH 8/RT, 200 mM KCl, 10 mM MgCl_2_, 1 mM DTT) using Zeba Spin Desalting Columns 7K (ThermoFisher Scientific #89883) before use. For complex assembly, core RNAP (E) and σ^N^ were mixed in reaction buffer and incubated for 10 min at 37°C to form Eσ^N^ followed by addition of the dhsU promoter fragment and another incubation for 10 min at 37°C to form the early-melted intermediate (RPem). C1 was added, and the reaction was incubated for 5 min at 37°C. Concentrations of each component in the final mix were 4 μM E, 4.8 μM σ^N^, 4.8 μM dhsU, and 5.6 μM C1. The Eσ^N^-dhsU-C1 complex was concentrated to 16 μM (using the concentration of E as a proxy for the complex) using Amicon Ultra 10K centrifugal filters (Merck Millipore #UFC501096). An ATP-detergent solution containing 40 mM ATP (Sigma #A2383) and 12 mM fluorinated fos-choline-8 (FC8F; Anatrace #F300F) in reaction buffer was prepared. ATP-detergent solution and Eσ^N^-dhsU-C1 were centrifuged at 11,000 xg for 10 min at 23°C before grid preparation.

#### Grid preparation

C-flat holey carbon grids (CF-1.2/1.3-4Au, EMS) were glow-discharged for 5 s at 25 mA and 0.3 mbar air atmosphere (Pelco easiGlow) before use. Sample application and vitrification was performed using a Vitrobot Mark IV (ThermoFisher Scientific) equilibrated to 37°C and 100% relative humidity in the blotting chamber. For each grid, the following procedure was performed: A glow-discharged grid was mounted in the blotting chamber of the Vitrobot instrument. 3.5 μL Eσ^N^-dhsU-C1 were equilibrated to 37°C on a thermoblock for at least 1 min. To start the reaction, 0.5 μL ATP-detergent solution was added. 3.5 μL of the reaction mix were immediately transferred to the grid, where reaction incubation was continued. Grids were blotted and plunged into liquid ethane with a total reaction time of 35-38 sec.

#### Cryo-EM data acquisition and processing

Grids were imaged using a 300 kV Titan Krios (ThermoFisher Scientific) equipped with a K3 camera (Gatan), a Cs corrector and a BioQuantum imaging filter (Gatan). Images were recorded using SerialEM ^57^ with a pixel size of 0.86 Å/px over a nominal defocus range of −0.8 μm to −2.2 μm and 20 eV energy filter slit width. A total of 17,199 gain-normalized movies were recorded in “super resolution” mode (K3 camera binning 0.5; image dimensions of 11,520 x 8,184 px; effective image pixel size of 0.43 Å/px) with 22 e^−^/px/s (at the camera) in dose-fractionation mode of 0.04 s over a 1.4 s exposure (35 frames) to give a total dose of 42 e^−^/Å^2^. Dose-fractionated movies were binned by a factor of 2 (resulting image pixel size of 0.86 Å/px), drift-corrected, summed, and dose-weighted using MotionCor2 ^58^. The contrast transfer function (CTF) was estimated for each summed image using the Patch CTF module in cryoSPARC v4.3.1 ^59^. Micrographs were curated to remove outliers in estimated CTF fit resolution (> 10 Å discarded), astigmatism (> 5000 Å discarded) and relative ice thickness (0.8 < x < 1.07 retained) resulting in a set of 16,649 images. Particles were picked using cryoSPARC Blob Picker (10,510,033 picks) retaining only picks with a normalized cross-correlation (NCC) score > 0.1 and a power score between 370 and 800 (NCC and power scores were scaled based on the entire set of picks). Particles were extracted from images with a box size of 448 px, Fourier-cropped to 128 px and subjected to two rounds of cryoSPARC 2D classification (number of classes, N=200) resulting in 1,341,601 particles. Duplicate particles were removed based on a center-to-center cutoff distance of 20 Å keeping the particles with higher NCC score. Images with less than 5 particles were manually inspected and excluded from further analysis (64 images/161 particles removed). Initial models were generated using cryoSPARC Ab initio Reconstruction ^59^ from a subset of 300,000 particles yielding reconstructions centered on RNAP (2 classes) or bEBP (2 classes) or “junk” (2 classes). Particles were further curated using two rounds of cryoSPARC Heterogeneous Refinement (N = 6) with ab initio classes serving as 3D references (2 RNAP classes, 2 x 2 junk classes). Particles contributing to RNAP or bEBP classes were combined and as above, duplicates and images with less than 5 particles were removed resulting in 1,060,259 particles (16,461 images). The particle stack was refined using cryoSPARC Non-Uniform (NU) Refinement with Defocus and Global CTF Refinement (tilt/trefoil) enabled ^60^ and then further processed using RELION v4.0.1 Bayesian Polishing ^61^. Polished particles were imported into cryoSPARC and subjected to NU Refinement with Defocus and Global CTF Refinement (tilt/trefoil/tetrafoil/spherical aberration) enabled, yielding a consensus reconstruction of 2.3 Å nominal resolution.

Initially, the consensus particle stack was subjected to cryoSPARC 3D classification (without masks) with various settings to explore the heterogeneity of the sample. To proceed, a 3D classification (N = 12, target resolution = 6 Å, random class initiation) using a focus mask around the bEBP and σ^N^-RI/bubble region (mask 1) was performed, providing improved particle assignments while yielding similar results to the unmasked classification. Classes were manually inspected and pooled yielding an “early-melted” class (295,087 particles), a “bEBP-bound” class (522,509 particles), and a “melted” class (133,297 particles) (Fig. 1C). Each class was subjected to NU Refinement ^60^.

The refined “melted” class (133,297 particles) was further divided by cryoSPARC 3D classification (N=10, target resolution=6 Å, no masks supplied, random class initiation). Resulting classes were manually inspected, pooled, and refined to yield maps for the “RPo” complex (45,631 particles, 2.8 Å) and “RPo+2A” complex (15,649 particles, 3.1 Å). For each complex, local refinement using a mask around σ^N^ (RPo-SigN mask) on signal subtracted particles was carried out. Locally refined maps were merged with the parent map using phenix.combine_focus_maps ^62^.

Further cryoSPARC 3D classification (N = 24, target resolution = 6 Å, random class initiation) of the refined “bEBP-bound” class (522,509 particles) was carried out using the focus mask around the bEBP and σ^N^-RI/bubble region (mask 1). Classes were manually pooled based on similarity and quality of σ^N^-RI density yielding three groups: “full helix” (74,144 particles), “one turn” (41,788 particles), and “two turns” (378,668 particles). A mask based on all orientations of the bEBP relative to RNAP (mask 2) was generated and used for signal subtraction of the “two turns” particle stack. Subtracted particles were subjected to local refinement using mask 2 followed by another local refinement using a tight mask around the bEBP from the first local refinement (mask 3). Using the tight mask around the bEBP (mask 3), cryoSPARC 3D classification (N = 10, target resolution = 4 Å, random class initiation) was performed yielding class pools “bEBP open” (118,561 particles, 3.1A) and “bEBP closed” (260,106 particles, 3.0 Å). For each class, the unsubtracted particles were aligned by NU Refinement to yield “RNAP” maps. To improve map quality in σ^N^ and the DNA, a mask around most parts of σ^N^ (excluding CBDs) and the σ^N^-bound DNA (SigN mask) was used to perform signal subtraction and local refinement yielding “SigN” maps. The bEBP and SigN maps were merged with the RNAP map using phenix.combine_focus_maps ^62^.

In a separate approach, the “bEBP-bound” class (522,509 particles) was again subjected to cryoSPARC 3D classification (N = 12, target resolution = 6 Å, forced hard classification, random class initiation) using a focus mask around the RNAP downstream channel (ds channel mask). While seven classes exhibited presence of DNA (295,571 particles; 56.6%), the remaining five classes contained density for an extended σ^N^-RI and σ^N^-RII (226,938 particles; 43.4%). The quality of the density for σ^N^-RI and σ^N^-RII varied considerably among the classes indicating flexibility, and therefore, only the two classes with the best resolved density were pooled and refined to yield the map “RII” (95,920 particles, 2.7 Å).

Local resolution and locally filtered maps were generated using cryoSPARC ^59^ or blocres/blocfilt ^63^. Most structural biology software was accessed through the SBGrid software package ^64^.

#### Cryo-EM analysis of C1-GS4-proteasome complexes

400 nM 20S-og and 800 nM C1-GS4 (final concentrations) were mixed in 40 mM Tris-HCl, pH 8/RT, 200 mM KCl, 10 mM MgCl_2_, 1 mM DTT, 0.5 mM ADP, 4 mM NaF, 1 mM AlCl_3_ and incubated for 5 min at 37°C. Grids (Ultrathin C on lacey carbon Cu400, TedPella #01824) were glow-discharged for 10 s at 10 mA and 0.3 mbar air atmosphere (Pelco easiGlow) before use. Vitrified samples on grids were prepared using a Vitrobot Mark IV (ThermoFisher Scientific) at 10°C and 100% relative humidity in the sample chamber. For each grid, 3.5 μL sample were applied to a continuous carbon grid and incubated for 30 s. Excess liquid was blotted away, and the grid was plunged into liquid ethane.

Data was collected on a 200 kV Talos Arctica instrument (ThermoFisher Scientific) equipped with a K2 camera (Gatan). Dose-fractionated movies were recorded in “counting” mode (image dimensions 3,838 x 3,710 px) using SerialEM ^57^ with a pixel size of 1.5 Å/px, 12s exposure, and 40 frames per movie. Dataset 1 was collected with a tilt angle of 20° over a nominal defocus range of −1 to −2.5 μm with a dose of 1.227 e^−^/Å^2^/frame and comprised 466 movies. Dataset 2 was collected with a tilt angle of 0° over a nominal defocus range of −0.8 to −3 μm with a dose of 1.12 e^−^/Å^2^/frame and comprised 503 movies. For each dataset, movies were drift-corrected, summed, and dose-weighted using MotionCor2 ^58^. Micrographs were further processed in cryoSPARC v4.3.1 ^59^ for each dataset separately. After CTF estimation using cryoSPARC’s PatchCTF module, particles were picked using circular blobs (BlobPicker; diameter 100-250 Å) and outliers were rejected based on NCC and power score resulting in 462,564 picks for dataset 1 and 475,471 picks for dataset 2. Particle images were extracted with a box size of 256 px and subjected to 2 rounds of 2D classification (120 classes, batch size 200) resulting in 291,510 particles for dataset 1 and 315,009 particles for dataset 2. For each dataset, ab initio classes were generated (3-6 classes, C7 symmetry) followed by a Non-Uniform Refinement (C7 symmetry) of all particles using the ab initio volume with the best quality as input. Duplicates with less than 50 Å center-to-center distance (based on 3D alignments) were removed, and particles were re-extracted with a box size of 288 px to recenter particle images resulting in 283,067 particles for dataset 1 and 308,092 particles for dataset 2. At this point, particle stacks from both datasets were merged, and new ab initio volumes were generated (6 classes, C7 symmetry). The merged particle stack was subjected to two rounds of Heterogeneous Refinement resulting in 437,119 particles. The particles were refined (Non-Uniform Refinement, C7 symmetry, per-particle defocus and global CTF refinement (tilt/trefoil) enabled) using the volume from the last Heterogenous Refinement run with the best quality as input yielding a reconstruction of 3.4 Å nominal FSC resolution. Duplicates with less than 20 Å center-to-center distance based on 3D alignments were removed and particle alignments were refined again (Non-Uniform Refinement, C7 symmetry, per-particle defocus and global CTF refinement (tilt/trefoil) enabled) yielding a reconstruction of 3.3 Å nominal FSC resolution from 434,340 particles. To improve density for the bEBP, the particles were subjected to 3D classification (10 classes, filter resolution = 6 Å, O-EM epochs = 4) using a mask around the bEBP. The resulting classes showed rotational variability of the bEBP density around the central particle axis, and single classes exhibited a hexagonal outline typical for the bEBP. One class with well-defined bEBP was chosen and subjected to local refinement using a solvent mask of the entire complex to obtain a reconstruction without symmetry constraints. The final volume was reconstructed from 43,660 particles at a nominal FSC resolution of 4.6 Å.

#### Proteolysis assays

RPem complexes were assembled by mixing E, σ^N^ (wild-type or variant), promoter DNA (dhsU or dhsU-CT), bEBP (C1 or C1-GS4), and 20S-og in buffer T [40 mM Tris-HCl, pH8/RT, 200 mM NaCl, 10 mM MgCl_2_, 1 mM DTT, 5% (v/v) glycerol]. At each addition step, the reaction was incubated for 10 min at 37°C to allow complex formation. Reactions were started with the addition of ATP (or alternative nucleotides or buffer as indicated) and incubated at 37°C for the indicated time. To stop the reaction, 9 μL reaction mix were directly mixed with an appropriate volume of SDS-PAGE loading dye and heated to 95°C for 5 min. Reaction components were separated by SDS-PAGE and stained with Coomassie for analysis. Final concentrations of the reaction components, if not stated otherwise, were 0.5 μM E, 0.5 μM σ^N^, 0.6 μM promoter DNA, 0.5 μM bEBP, 0.6 μM 20S-og, and 5 mM ATP.

#### *In vitro* transcription assays

Reactions were performed in buffer R (40 mM Tris-HCl, pH8/RT, 200 mM KCl, 10 mM MgCl_2_, 1 mM DTT, 5 μg/ml BSA) and assembled as follows: E and σ^N^ were mixed and incubated for 10 min at 37°C to form Eα^N^. Promoter fragment dhsU was added, and the reaction was incubated for 10 min at 37°C to form RPem. bEBP (C1 or C1-GS4) was added and the reaction was incubated for up to 5 min at 37°C. Reactions were started with addition of NTP mix (ATP, GTP, CTP, UTP, and α-^32^P-UTP) and incubated at 37°C for 15 min before stopping the reaction by adding an equal volume of 2x STOP buffer [0.5x TBE, 8 M urea, 30 mM EDTA, 0.05% (w/v) bromophenol blue, 0.05% xylene cyanol]. Stopped reactions were heated at 95°C for 5 min before separation on a 20% (1:29 acrylamide:bis-acrylamide) 1x TBE-urea gel. Bands were visualized by autoradiography. Final concentrations in the reactions were: 80 nM E, 200 nM σ^N^, 10 nM promoter DNA, 1 μM bEBP, 1.1 μM 20S-og, 5 mM ATP, 0.5 mM GTP, 0.5 mM CTP, 0.05 mM UTP, and 0.1 μCi/μl α-^32^P-UTP unless noted otherwise.

#### Model building and refinement

Initial models were derived from PDBs 8F1K ^21^, 6GH5 ^42^ and 4LZZ ^65^. Models were manually fit into the cryo-EM density maps using Chimera (Pettersen et al., 2004) and rigid-body refined using phenix.real_space_refine ^66^. Models were inspected and modified in Coot ^67^ or ISOLDE ^68^. Models were finalized by all-atom and B-factor refinement with Ramachandran and secondary structure restraints using phenix.real_space_refine. Maps and models were visualized in ChimeraX ^69^.

#### Sequence alignments

Protein sequences were aligned using Clustal Omega ^70^. Sequence motifs were generated using WebLogo3 ^71^. The conserved net charge U at position n of the aligned sequences was calculated as follows

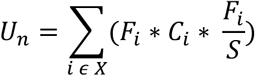

where X is the set of all possible amino acids, i.e., X = {Ala, Gly, Val, Leu, Met, Arg, Cys, Phe, Tyr, Trp, …}, and

F_i_ = frequency of residue i

C_i_ = net charge of residue i at physiological pH

S = total number of sequences in the alignment

Net charges at physiological pH were assumed as R/K = +1, D/E = −1 and any other = 0.

## Data and materials availability

Model and map files were deposited at the Protein Data Bank (PDB) and Electron Microscopy Data Bank (EMDB), respectively: RPi1^open^ (PDB 9MSE, EMD-48586, EMD-48580, EMD-48581, EMD-48582), RPi1^closed^ (PDB 9MSF, EMD-48587, EMD-48583, EMD-48584, EMD-48585), RPo (PDB 9MSH, EMD-48589, EMD-48576, EMD-48577), RPo+2A (PDB 9MSJ, EMD-48590, EMD-48578, EMD-48579), RII (PDB 9MSG, EMD-48588), and 20Sog+C1-GS4+ADP-AlFx (EMD-48574). All other data are available in the manuscript or the supplementary materials. Reasonable requests for materials should be submitted to the Lead Contact, Seth A. Darst.

## Extended Data

**Extended Data Fig. 1:**
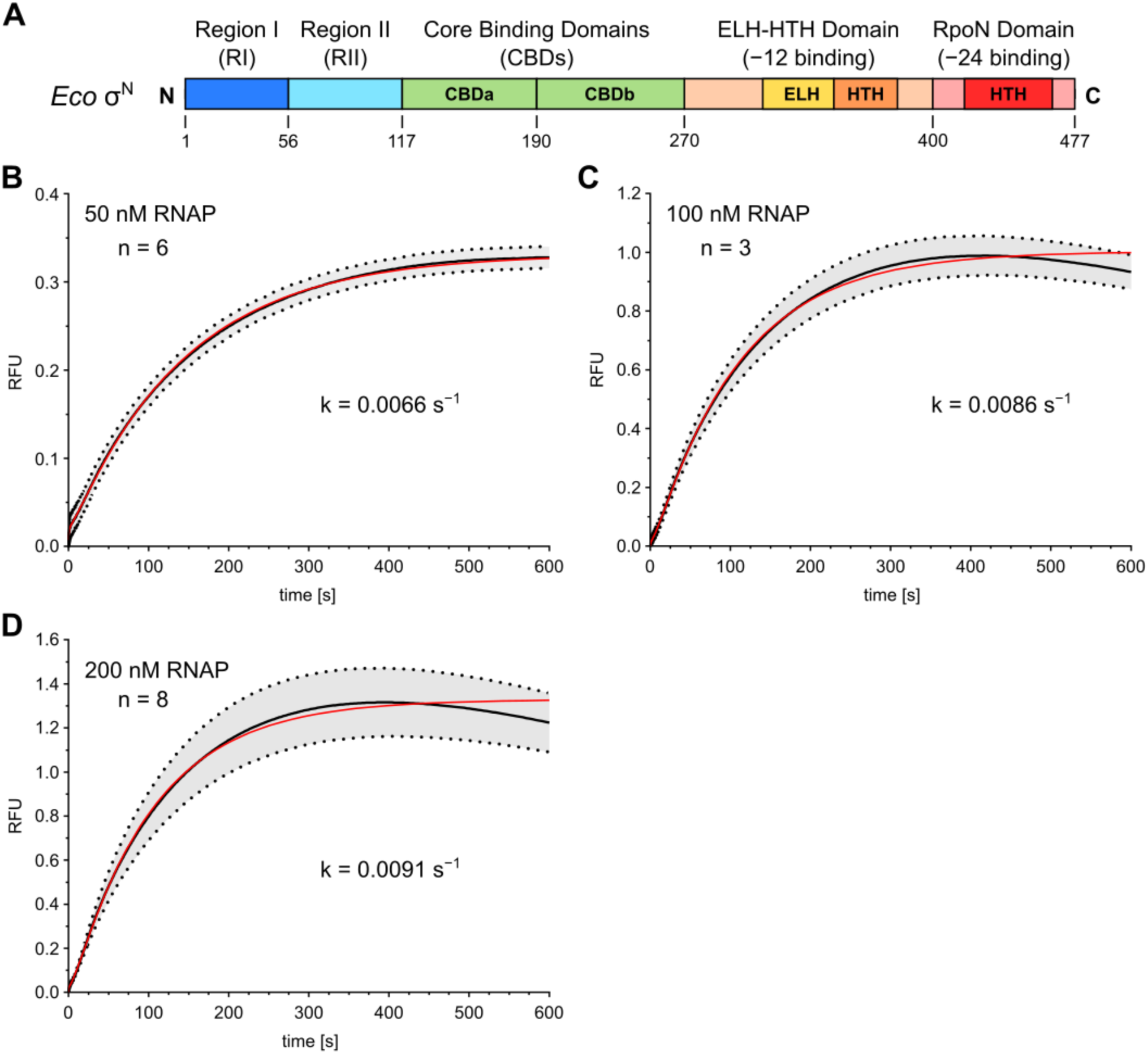
RPo formation kinetics of *Eco* Eσ^N^ on the *Aae* dhsU promoter in presence of a constitutively active *Aae* NtrC1 variant (residues 121-387). (A) Schematic of *E. coli* (*Eco*) σ^N^ domain architecture showing N-terminal region I (RI; 1-56), region II (RII; 57-117), the two core binding domains (CBDs; CBDa: 118-190, CDBb: 191-270), the extra-long helix (ELH) helix-turn-helix (HTH) domain (270-400), and the C-terminal RpoN domain (401-477). (B-E) Stopped flow measurements of RPo formation on the labeled dhsU+2T-Cy3 promoter fragment at (B) 50 nM (n = 6), (C) 100 nM (n = 3), and (D) 200 nM (n = 8) complex in presence of bEBP and ATP. The black line shows the average of n individual mixing events. Error bands (gray, dotted lines) represent the standard deviation. Data fitting (red trace) was performed as described in the methods section. Data in (B) is the same as shown in Fig. 1B.

**Extended Data Fig. 2:**
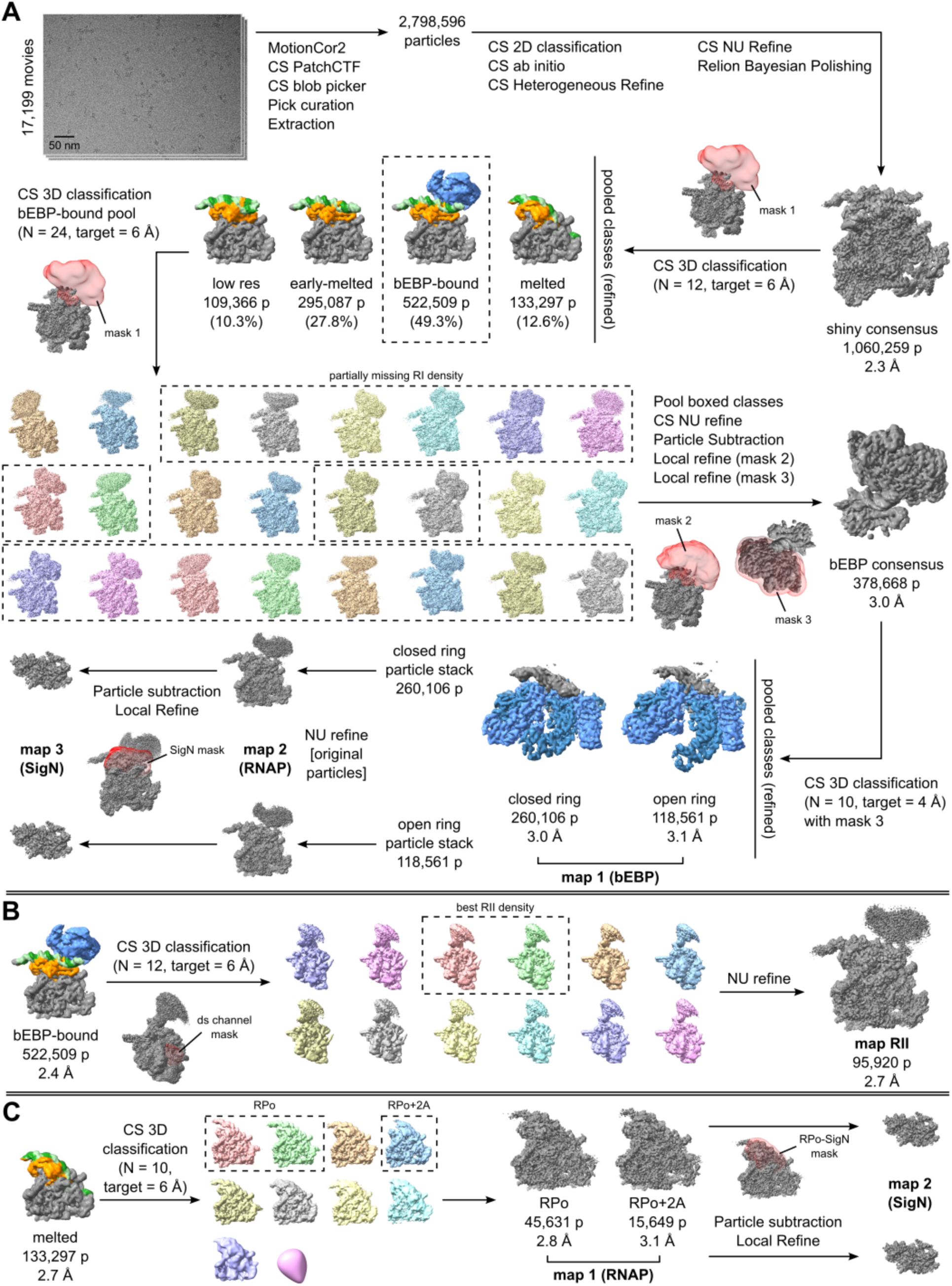
Overview of the cryo-EM data processing. (A) Initial processing of raw movies to a consensus particle stack yielding a “shiny consensus” map. The consensus particle stack was subjected to masked 3D classification in cryoSPARC (CS). The resulting group of bEBP-bound complexes was further processed by several rounds of 3D classification and local refinements yielding maps for the “open” and “closed” ring states. (B) Subclassification of bEBP-bound complexes with ds channel mask. (C) Subclassification of melted complexes. See methods section for details.

**Extended Data Fig. 3:**
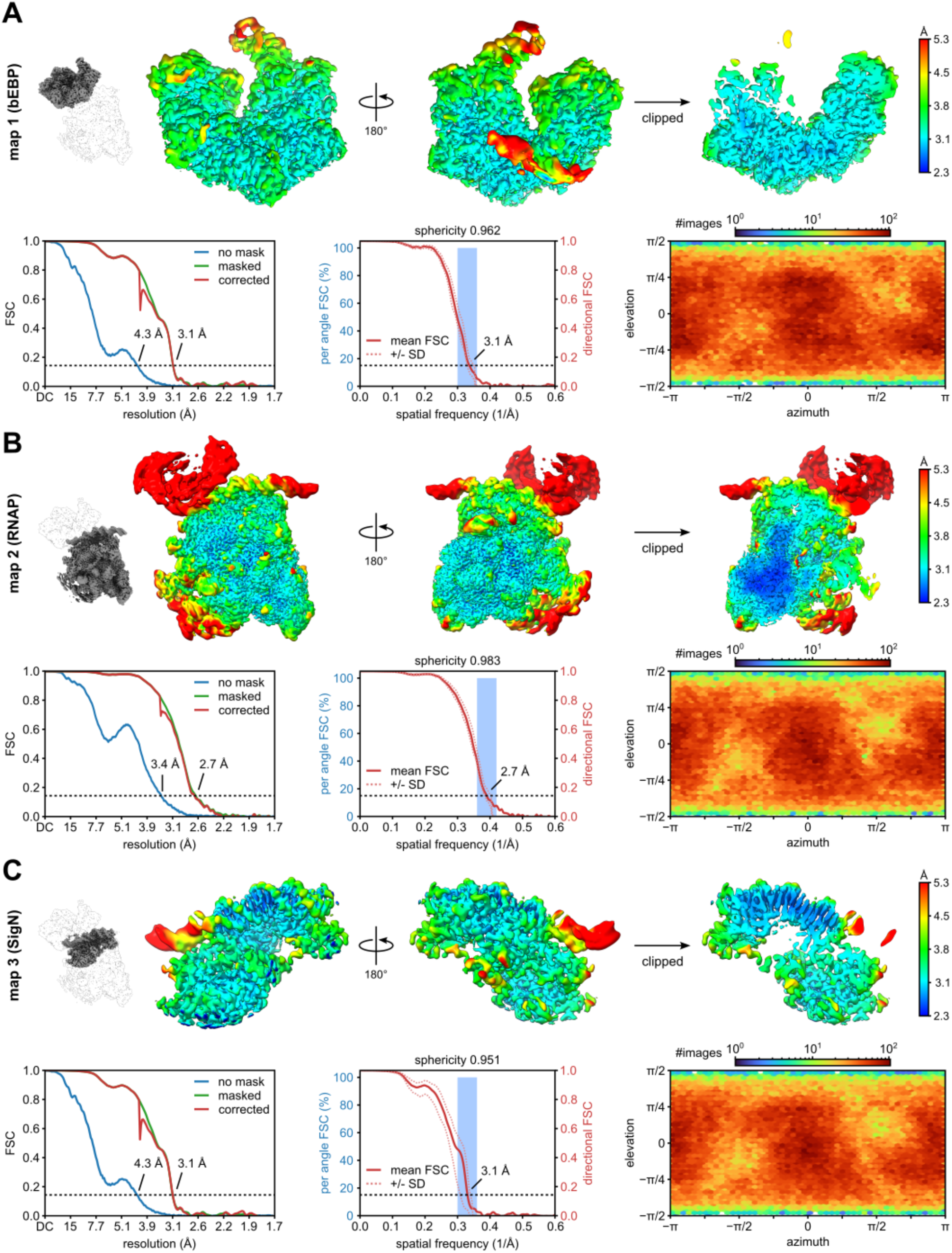
Local resolution, FSC, and particle orientation of the open ring state (RPi1^open^). (A) bEBP focused map (map 1). (B) RNAP map (map 2). (C) σ^N^ focused map (map 3). The small inset depicts the map location in the full complex. For each map, views of the locally filtered map are shown in the top row and colored according to local resolution. Bottom left panel: FSC curves calculated without mask (blue), masked (green), and corrected by phase randomization and noise substitution (red). Middle panel: 3D-FSC plot showing mean FSC curve (solid line) and FSC curves +/− 1 standard deviation (dotted lines). Blue histogram bars depict frequency of FSC estimates from a total of FSC calculations at 100 angled cones (each cone 20°). Bottom right panel: Heat map of particle orientation assignments.

**Extended Data Fig. 4:**
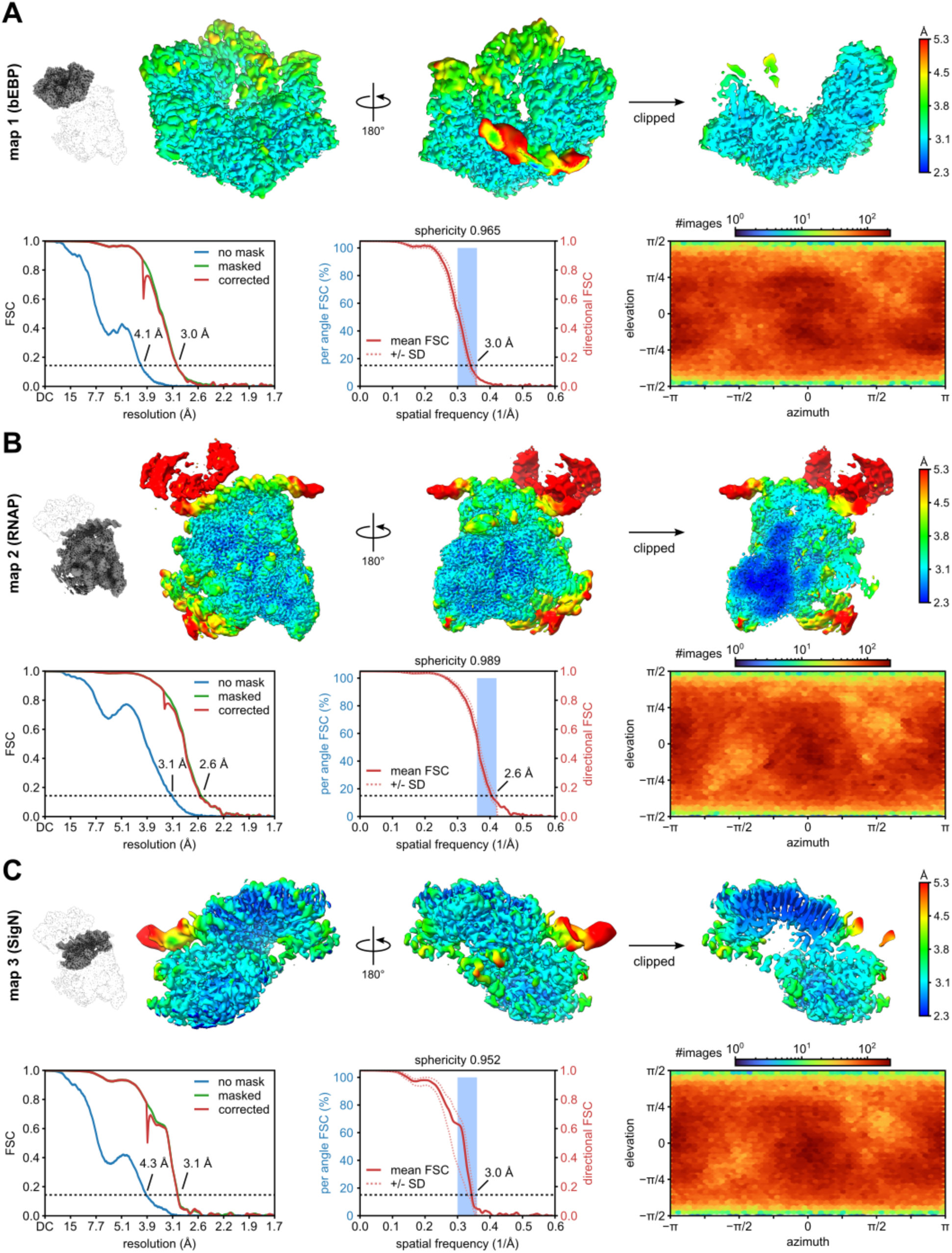
Local resolution, FSC, and particle orientation of the closed ring state (RPi1^closed^). (A) bEBP focused map (map 1). (B) RNAP map (map 2). (C) σ^N^ focused map (map 3). The small inset depicts the map location in the full complex. For each map, views of the locally filtered map are shown in the top row and colored according to local resolution. Bottom left panel: FSC curves calculated without mask (blue), masked (green), and corrected by phase randomization and noise substitution (red). Middle panel: 3D-FSC plot showing mean FSC curve (solid line) and FSC curves +/− 1 standard deviation (dotted lines). Blue histogram bars depict frequency of FSC estimates from a total of FSC calculations at 100 angled cones (each cone 20°). Bottom right panel: Heat map of particle orientation assignments.

**Extended Data Fig. 5:**
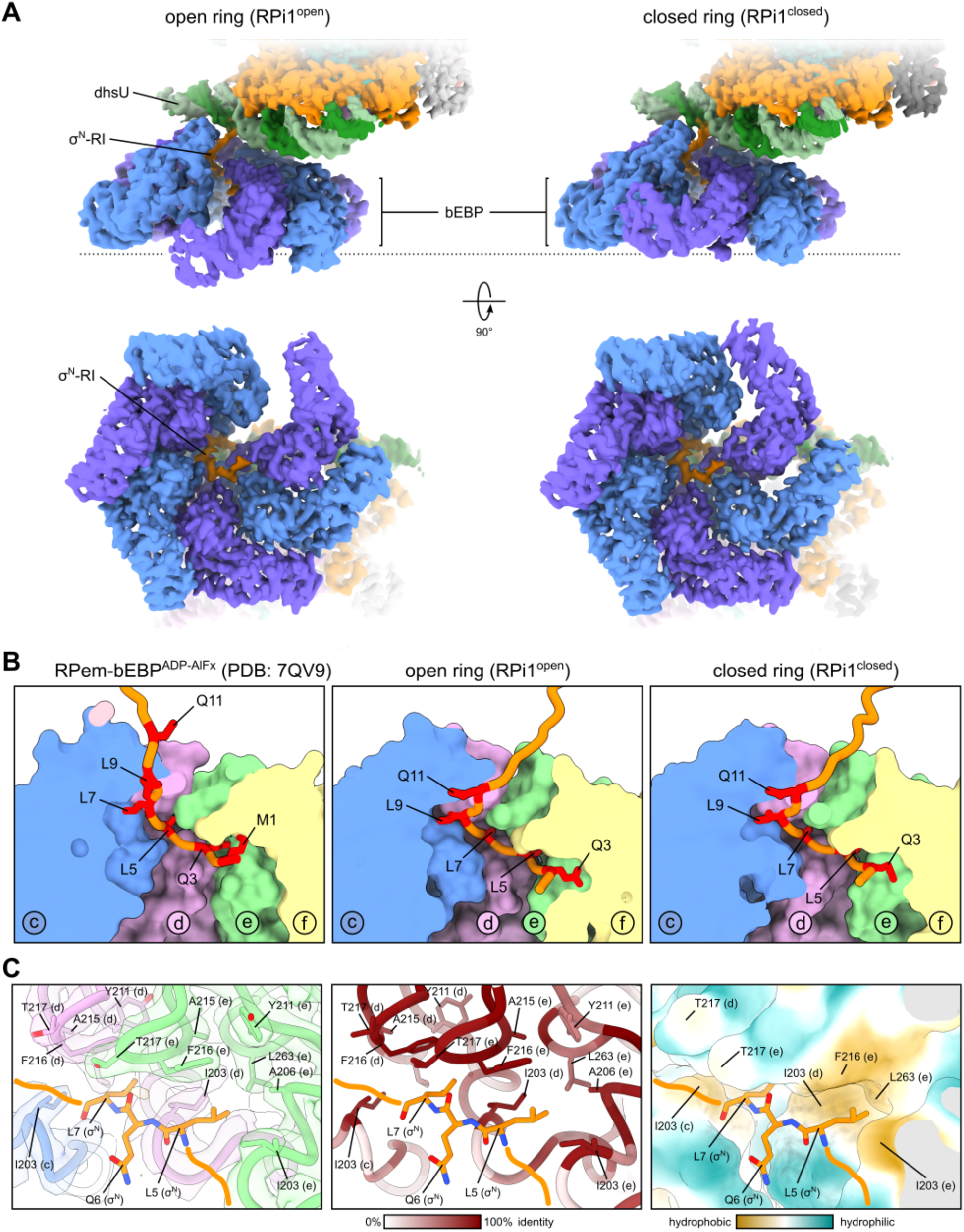
Comparison of bEBP ring state maps, σ^N^-RI translocation and pore pockets. (A) Unsharpened map of the open ring state (left) and closed ring state (right) colored according to subunit. Side (top row) and bottom views (bottom row) of the ring reveal clear density of σ^N^-RI inside the pore. (B) Comparison of the σ^N^-RI/bEBP interaction between ADP-AlFx bound bEBP (left), open ring (middle), and closed ring state (right). Every second residue in σ^N^-RI starting with M1 up to Q11 is highlighted in red. M1 was not modeled for the open and closed ring structures. bEBP subunits colored as in Fig. 2B-D. (C) Pore pockets in bEBP open complex with map density shown as transparent surfaces and colored according to subunit (left), conservation (middle; percent identity), and lipophilicity (right; calculated in ChimeraX). Sequences for the conservation coloring alignment were obtained via the HMMER webserver (phmmer; search against rp15 database) using the *A. aeolicus* NtrC1 sequence (UniProt ID O67198) as query. Sequences with an e-value > 10^−40^, smaller than 380 residues, and larger than 500 residues were excluded, and the result was clustered using CD-HIT (options -n 3 -c 0.5 -T 4) yielding 1484 representative sequences, which were aligned using Clustal Omega.

**Extended Data Fig. 6:**
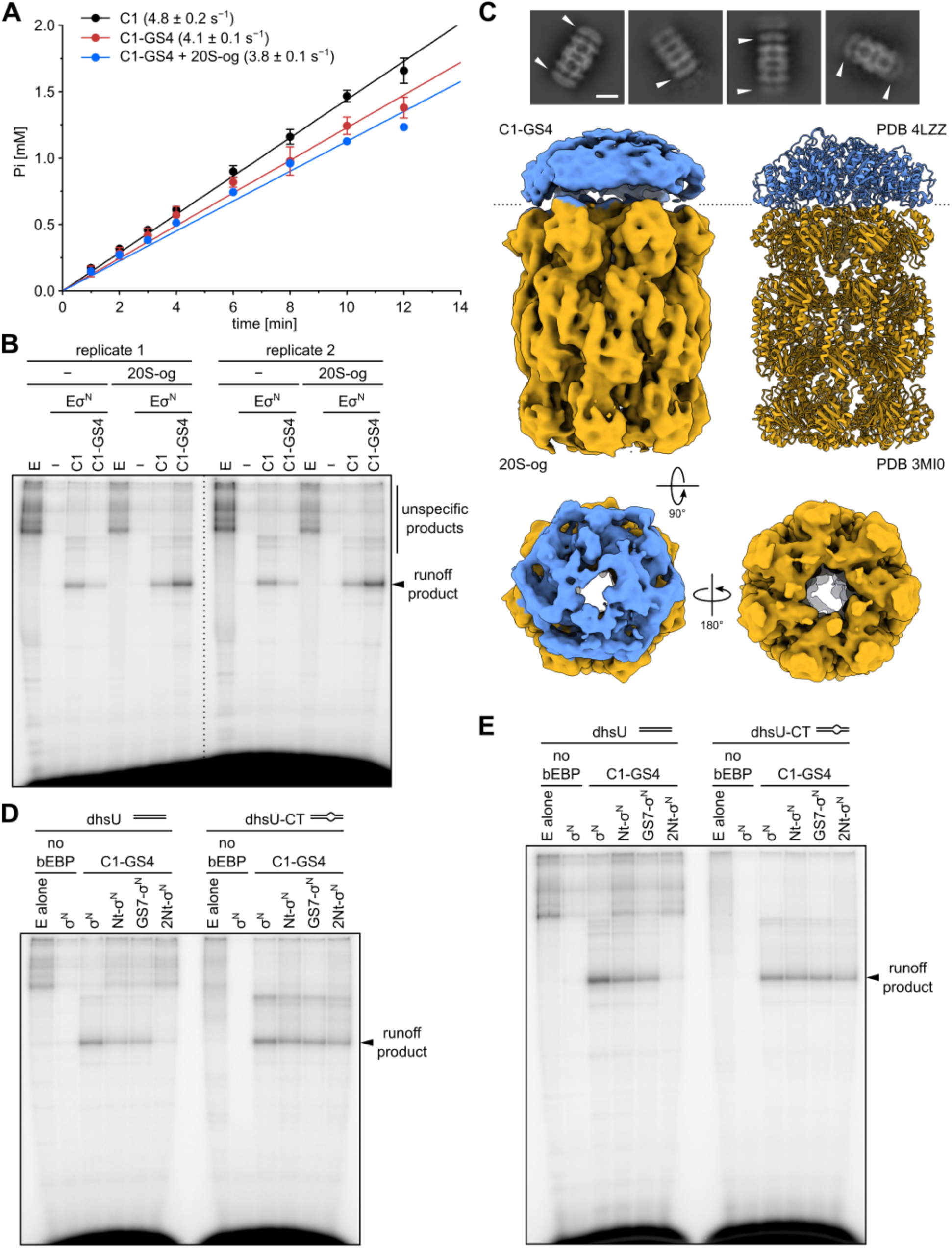
Validation of proteolysis assay. (A) C1-GS4 in presence or absence of the proteasome (20S-og) exhibits a similar ATPase activity to C1. Reactions were carried out at 37 °C and contained 0.5 μM bEBP (C1 or C1-GS4), 0.75 μM Mtb 20S-og, and 5 mM ATP. Data points show the mean and standard deviation of three replicates. (B) In vitro transcription from the dhsU promoter fragment with core RNAP (E), holoenzyme (Eσ^N^), and no bEBP, C1 or C1-GS4 in absence or presence of 20S-og. Reactions contained 10 nM promoter DNA, 80 nM E, 200 nM σ^N^, 1 μM bEBP, 1.1 μM 20S-og, 5 mM ATP, 0.5 mM GTP, 0.5 mM CTP, 0.05 mM UTP, and 0.1 μCi/ul α-^32^P-UTP Two replicates of individual reactions are shown. (C) Cryo-EM imaging of the C1-GS4/proteasome complex. Exemplary side view 2D classes show capping of the proteasome particle by the bEBP variant C1-GS4 (top panel). White arrows indicate bEBP locations. Scale bar corresponds to 100 Å. The 3D reconstruction of the capped proteasome particle exhibits the hexagonal symmetry of the bEBP and the heptagonal symmetry of the proteasome. (D-E) In vitro transcription of σ^N^ variants from the dhsU or dhsU-CT promoter fragment. Reaction conditions as in (B). Each panel shows the result of an individual experiment.

**Extended Data Fig. 7:**
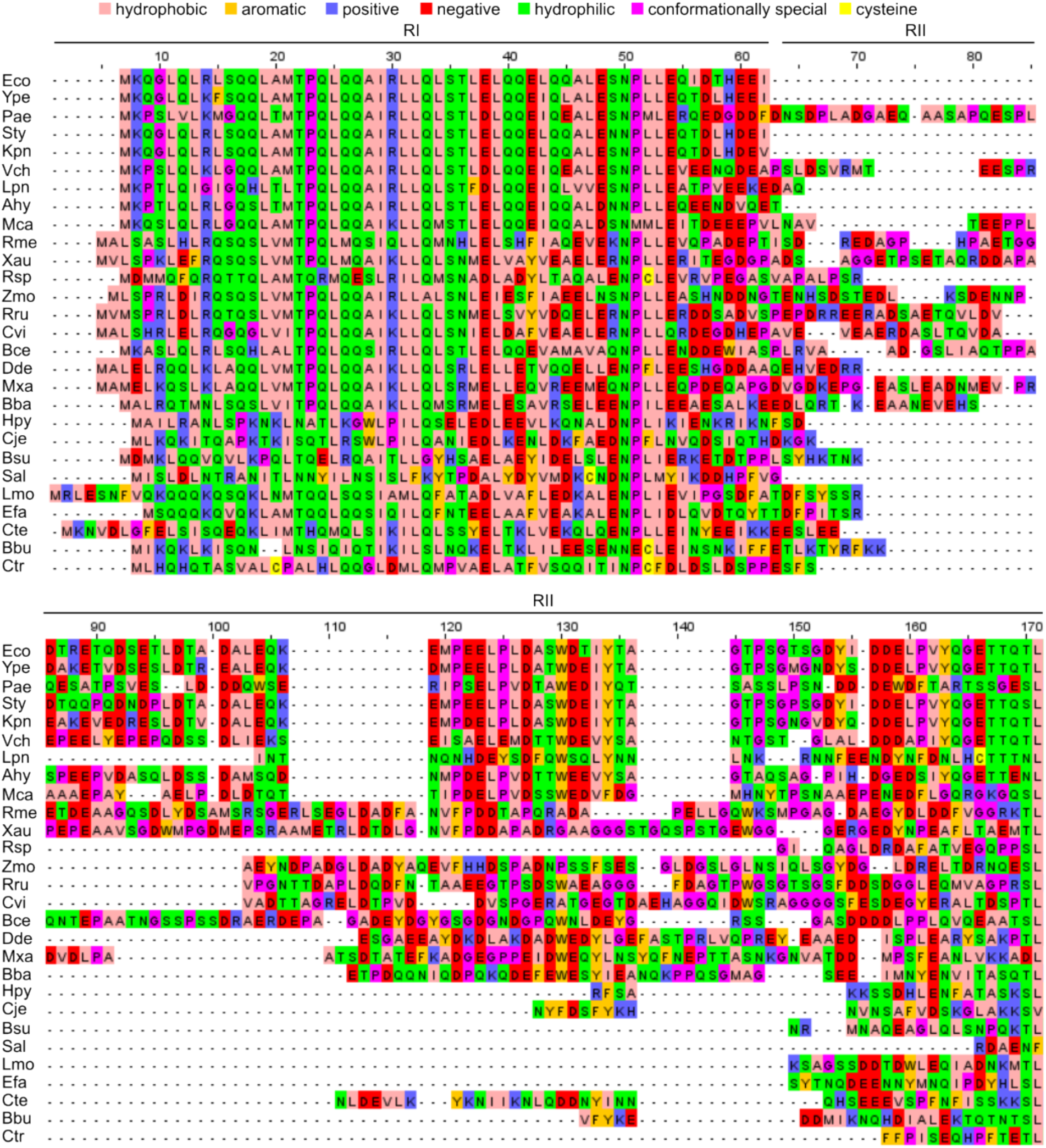
Alignment of σ^N^ protein sequences from exemplary species across the bacterial domain. The first 170 positions of the alignment encompassing RI and RII are shown. Residues are colored according to physicochemical properties: hydrophobic (salmon; I, L, V, A, M), aromatic (gold; F, W, Y), positive charge (blue; K, R, H), negative charge (red; D, E), hydrophilic (green; S, T, N, Q), special conformation (magenta; P, G), and cysteine (yellow). Regions of σ^N^ are indicated above the alignment. Eco = Escherichia coli (P24255), Ype = Yersinia pestis (A0A3N4AWI2), Pae = Peudomonas aeruginosa (P49988), Sty = Salmonella typhimurium (P26979), Kpn = Klebsiella pneumoniae (A0A0H3H3L1), Vch = Vibrio cholerae (C3LRJ2), Lpn = Legionella pneumophila (Q5WZ64), Ahy = Aeromonas hydrophila (A0KQ11), Mca = Methylococcus capsulatus (Q60AV3), Rme = Rhizobium meliloti (P17263), Xau = Xanthobacter autotrophicus (Q8RM50), Rsp = Rhodobacter sphaeroides (Q01194), Zmo = Zymomonas mobilis (Q5NQV6), Rru = Rhodospirillum rubrum (Q2RYI0), Cvi = Caulobacter vibrioides (Q03408), Bce = Burkholderia cenocepacia (B4EAS4), Dde = Desulfovibrio desulfuricans (B8IZF4), Mxa = Myxococcus xanthus (Q1DDF1), Bba = Bdellovibrio bacteriovorus (Q6MPK9), Hpy = Helicobacter pylori (P56143), Cje = Campylobacter jejuni (Q0PAK4), Bsu = Bacillus subtilis (P24219), Sal = Salinicoccus alkaliphilus (A0A1M7ILC4), Lmo = Listeria monocytogenes (Q7AP50), Efa = Enterococcus faecalis (Q837Q1), Cte = Clostridium tetani (Q898R6), Bbu = Borrelia burgdorferi (O51406), Ctr = Chlamydia trachomatis (A0A0H2X1C4). Parentheses contain UniProt accession numbers.

**Extended Data Fig 8:**
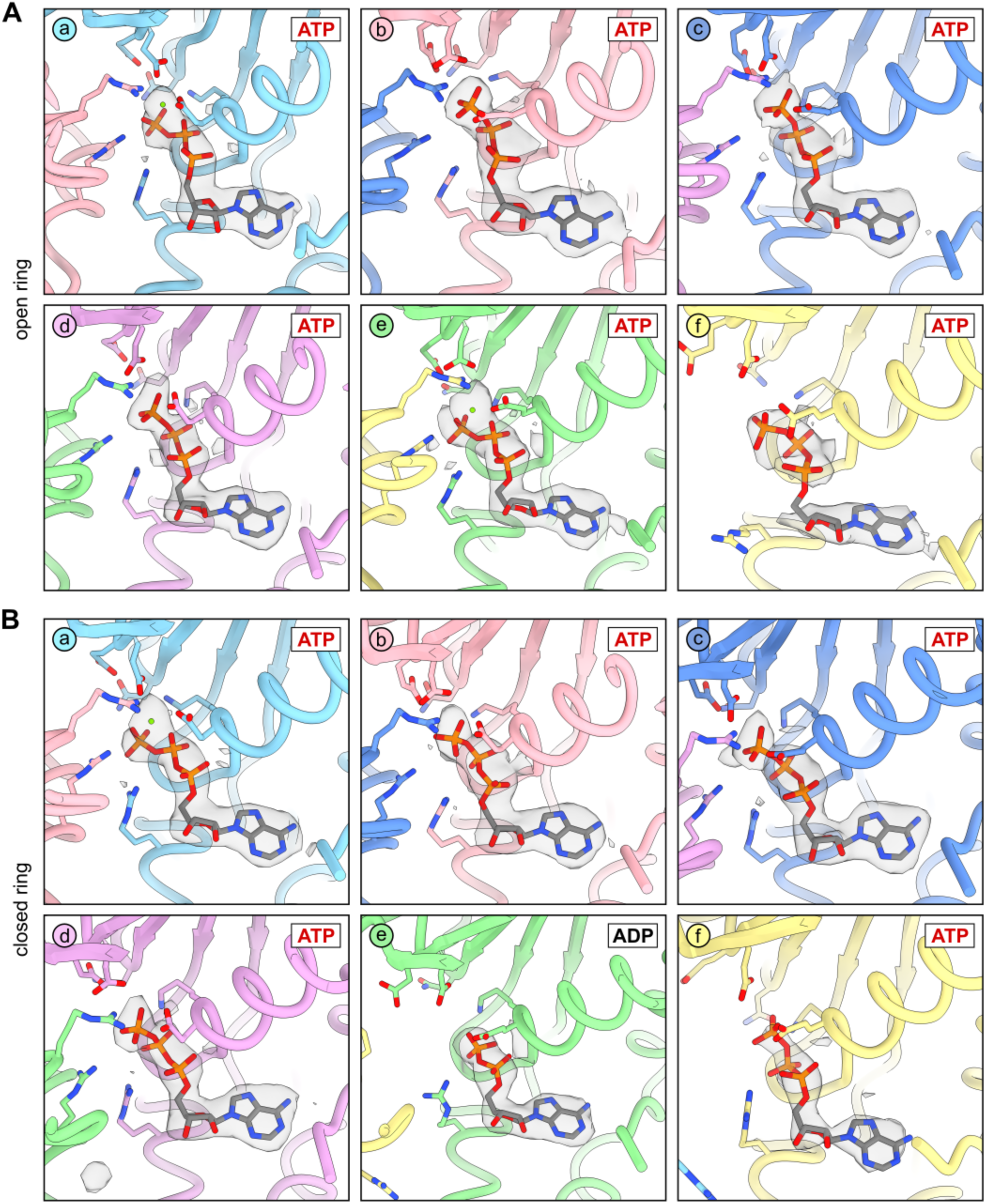
Nucleotide binding sites in the bEBP. Nucleotide binding pockets in the open (A) and closed ring state (B) of RPi1. Overlayed grey transparent surface represents the map density of the respective sharpened map. Relevant active site residues are shown in stick representation (see also main figures).

**Extended Data Fig. 9:**
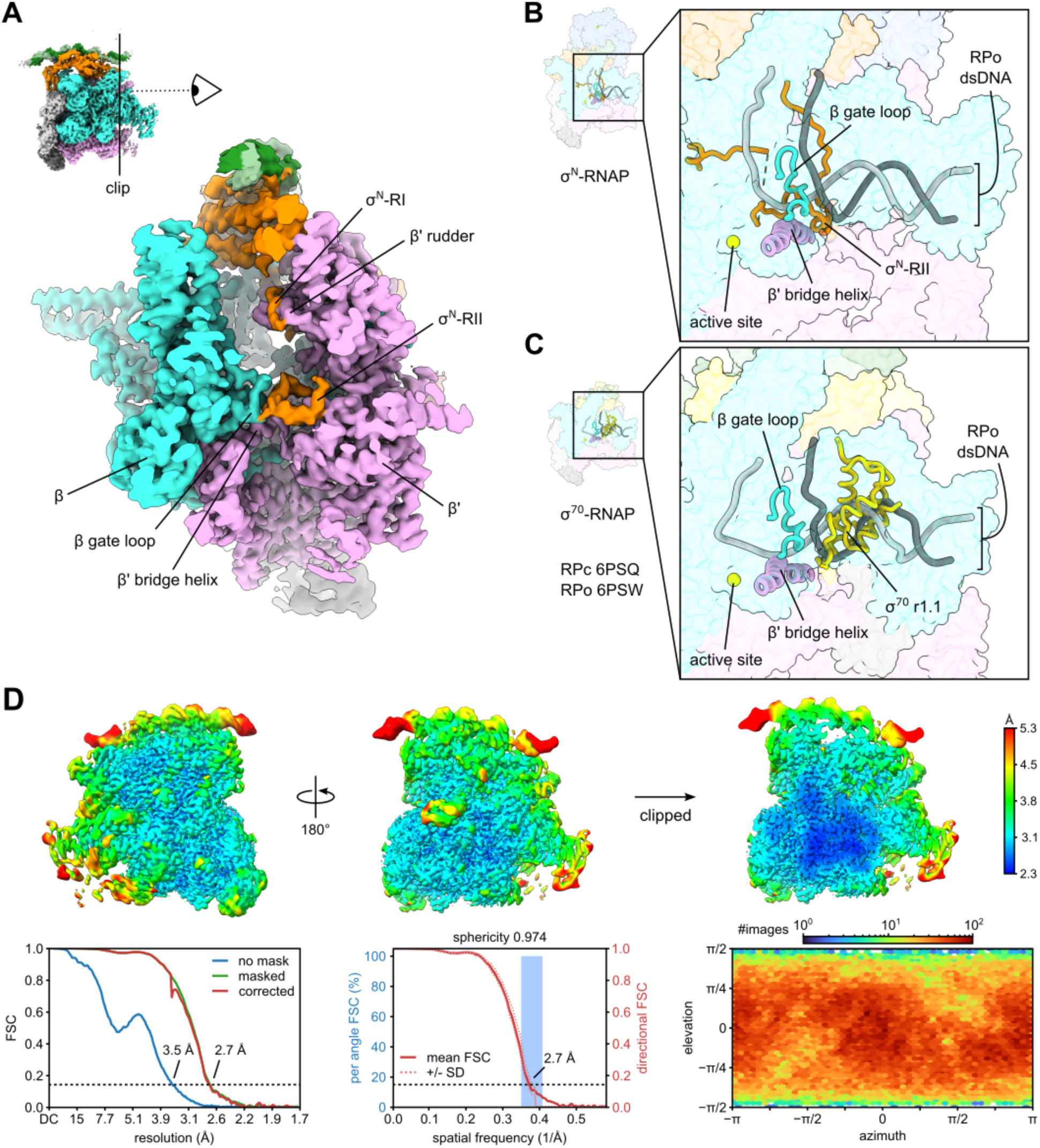
Binding of σ^N^-RII in the downstream channel of RNAP. (A) View along the axis of the downstream channel reveals density for σ^N^-RII winding between the β gate loop and the β’ bridge helix. Unsharpened map colored according to subunits; clipped for visual clarity. Note that the bEBP density is below the map threshold due to the rotational variability of the bEBP (see main text and methods for details). (B) Close-up view of the steric clash between RII and the downstream duplex DNA path in RPo. Cartoon of downstream DNA duplex from RPo overlayed in gray. Active site Mg^2+^ colored in yellow. (C) σ^70^ region 1.1 occupies downstream channel in RPc (PDB 6PSQ and 6PSW). (D) Views of the locally filtered map are colored according to local resolution. Bottom left panel: FSC curves calculated without mask (blue), masked (green), and corrected by phase randomization and noise substitution (red). Middle panel: 3D-FSC plot showing mean FSC curve (solid line) and FSC curves +/− 1 standard deviation (dotted lines). Blue histogram bars depict frequency of FSC estimates from a total of FSC calculations at 100 angled cones (each cone 20°). Bottom right panel: Heat map of particle orientation assignments.

**Extended Data Fig. 10:**
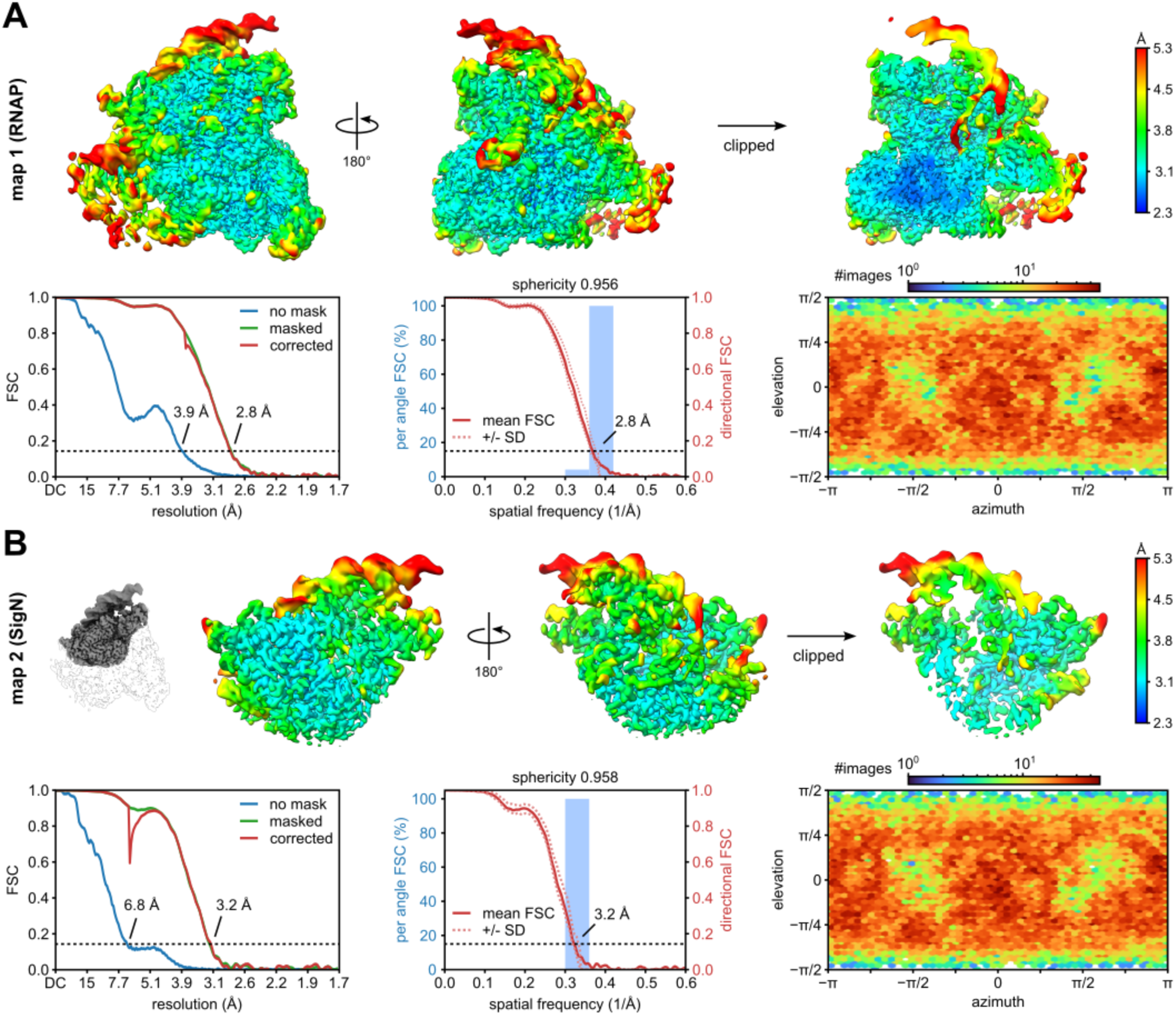
Local resolution, FSC, and particle orientation of the RPo state. (A) RNAP map (map 1). (B) σ^N^ focused map (map 2). The small inset depicts the map location in the full complex. For each map, views of the locally filtered map are shown in the top row and colored according to local resolution. Bottom left panel: FSC curves calculated without mask (blue), masked (green), and corrected by phase randomization and noise substitution (red). Middle panel: 3D-FSC plot showing mean FSC curve (solid line) and FSC curves +/− 1 standard deviation (dotted lines). Blue histogram bars depict frequency of FSC estimates from a total of FSC calculations at 100 angled cones (each cone 20°). Bottom right panel: Heat map of particle orientation assignments.

**Extended Data Fig. 11:**
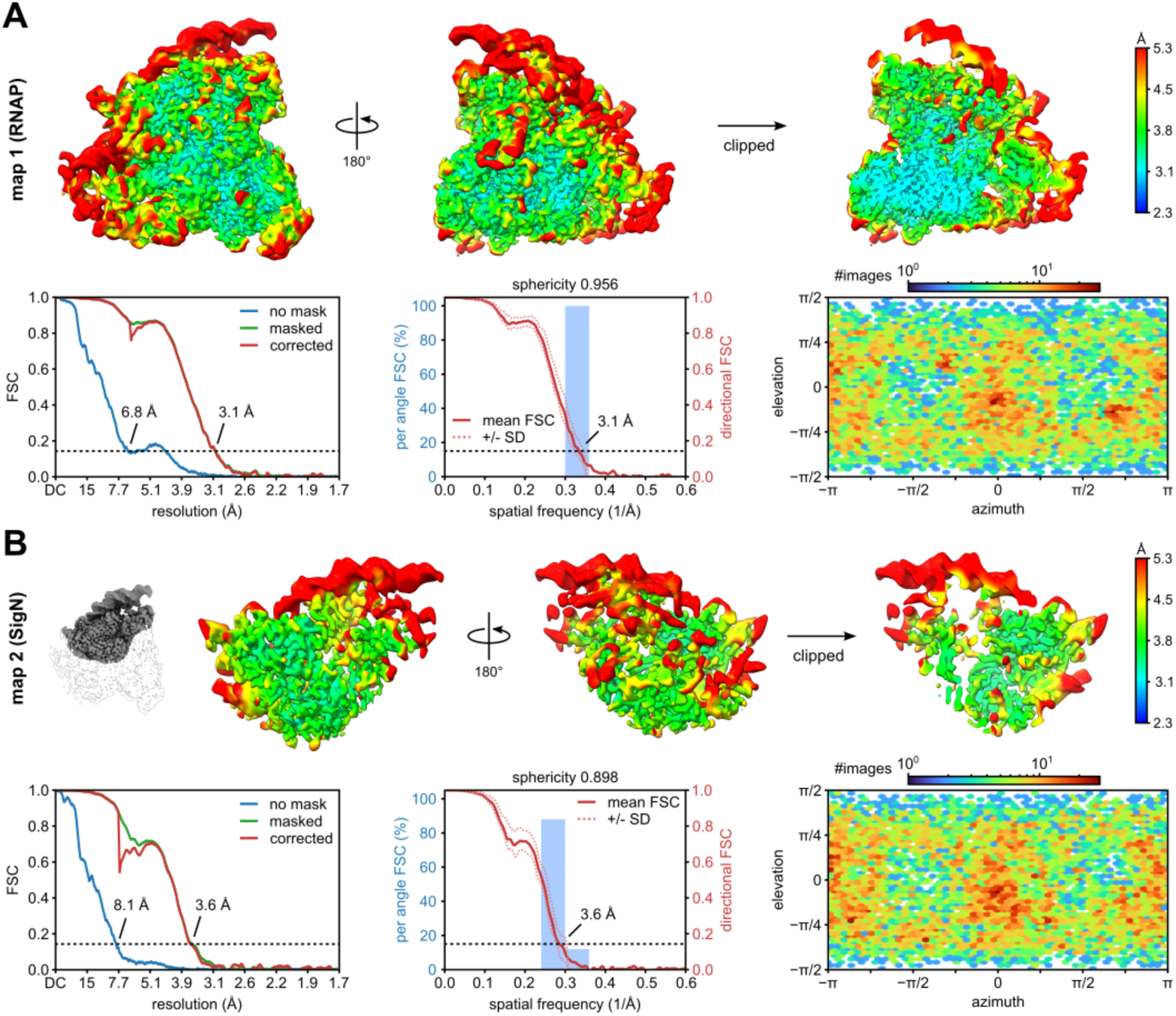
Local resolution, FSC, and particle orientation of the RPo+2A state. (A) RNAP map (map 1). (B) σ^N^ focused map (map 2). The small inset depicts the map location in the full complex. For each map, views of the locally filtered map are shown in the top row and colored according to local resolution. Bottom left panel: FSC curves calculated without mask (blue), masked (green), and corrected by phase randomization and noise substitution (red). Middle panel: 3D-FSC plot based on half maps showing mean FSC curve (solid line) and FSC curves +/− 1 standard deviation (dotted lines). Blue histogram bars depict frequency of FSC estimates from a total of FSC calculations at 100 angled cones (each cone 20°). Bottom right panel: Heat map of particle orientation assignments.

**Extended Data Fig. 12:**
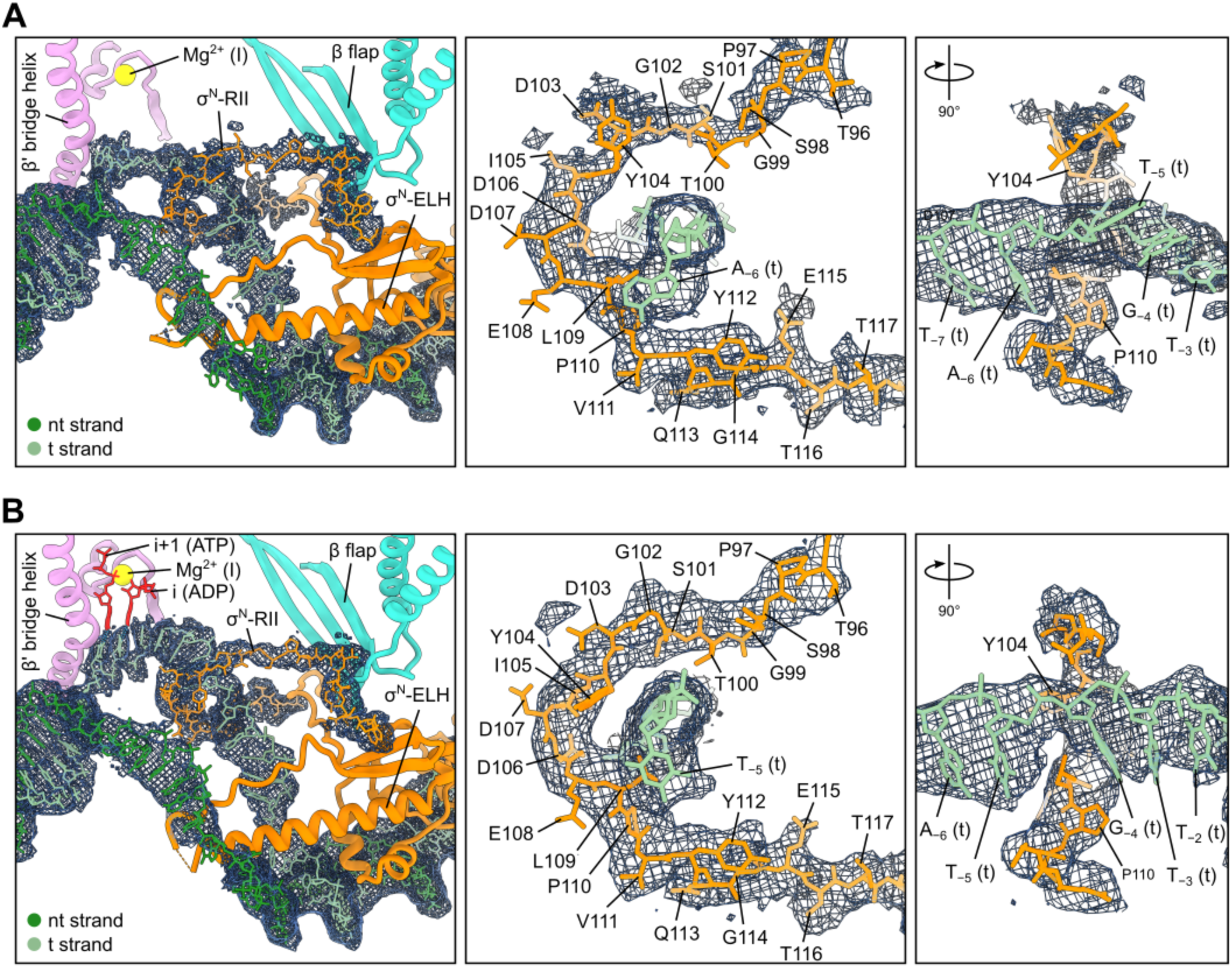
Entanglement of σ^N^-RII and the template strand. (A) in RPo. (B) in RPo+2A. Density of the locally filtered map around σ^N^-RII and the DNA is shown as meshed surface.

## References

1. Gruber, T.M., and Gross, C.A. (2003). Multiple Sigma Subunits and the Partitioning of Bacterial Transcription Space. Annu Rev Microbiol 57, 441–466. 10.1146/annurev.micro.57.030502.090913.

2. Lonetto, M., Gribskov, M., and Gross, C.A. (1992). The sigma 70 family: sequence conservation and evolutionary relationships. J Bacteriol 174, 3843–3849. 10.1128/jb.174.12.3843-3849.1992.

3. Studholme, D.J., and Dixon, R. (2003). Domain Architectures of σ54-Dependent Transcriptional Activators. J Bacteriol 185, 1757–1767. 10.1128/JB.185.6.1757-1767.2003.

4. Buck, M., Gallegos, M.-T., Studholme, D.J., Guo, Y., and Gralla, J.D. (2000). The Bacterial Enhancer-Dependent ς^54^ (ς ^N^) Transcription Factor. J Bacteriol 182, 4129–4136. 10.1128/JB.182.15.4129-4136.2000.

5. Correa, N.E., Lauriano, C.M., McGee, R., and Klose, K.E. (2000). Phosphorylation of the flagellar regulatory protein FlrC is necessary for Vibrio cholerae motility and enhanced colonization. Mol Microbiol 35, 743–755. 10.1046/j.1365-2958.2000.01745.x.

6. Feldman, M., Bryan, R., Rajan, S., Scheffler, L., Brunnert, S., Tang, H., and Prince, A. (1998). Role of Flagella in Pathogenesis of Pseudomonas aeruginosa Pulmonary Infection. Infect Immun 66, 43–51. 10.1128/IAI.66.1.43-51.1998.

7. Hathroubi, S., Zerebinski, J., and Ottemann, K.M. (2018). Helicobacter pylori Biofilm Involves a Multigene Stress-Biased Response, Including a Structural Role for Flagella. mBio 9, e01973–18. 10.1128/mBio.01973-18.

8. Soules, K.R., LaBrie, S.D., May, B.H., and Hefty, P.S. (2020). Sigma 54-Regulated Transcription Is Associated with Membrane Reorganization and Type III Secretion Effectors during Conversion to Infectious Forms of Chlamydia trachomatis. mBio 11, e01725–20. 10.1128/mBio.01725-20.

9. Fisher, M.A., Grimm, D., Henion, A.K., Elias, A.F., Stewart, P.E., Rosa, P.A., and Gherardini, F.C. (2005). Borrelia burgdorferi sigma 54 is required for mammalian infection and vector transmission but not for tick colonization. Proceedings of the National Academy of Sciences 102, 5162–5167. 10.1073/pnas.0408536102.

10. Hauser, F., Pessi, G., Friberg, M., Weber, C., Rusca, N., Lindemann, A., Fischer, H.-M., and Hennecke, H. (2007). Dissection of the Bradyrhizobium japonicum NifA+σ54 regulon, and identification of a ferredoxin gene (fdxN) for symbiotic nitrogen fixation. Molecular Genetics and Genomics 278, 255–271. 10.1007/s00438-007-0246-9.

11. Bush, M., and Dixon, R. (2012). The Role of Bacterial Enhancer Binding Proteins as Specialized Activators of σ ^54^ -Dependent Transcription. Microbiology and Molecular Biology Reviews 76, 497–529. 10.1128/MMBR.00006-12.

12. Wang, J.T., Syed, A., Hsieh, M., and Gralla, J.D. (1995). Converting Escherichia coli RNA Polymerase into an Enhancer-Responsive Enzyme: Role of an NH2-Terminal Leucine Patch in sigma54. Science (1979) 270, 992–994. 10.1126/science.270.5238.992.

13. Syed, A., and Gralla, J.D. (1997). Isolation and properties of enhancer-bypass mutants of sigma 54. Mol Microbiol 23, 987–995. 10.1046/j.1365-2958.1997.2851651.x.

14. Wang, J.T., Syed, A., and Gralla, J.D. (1997). Multiple pathways to bypass the enhancer requirement of sigma 54 RNA polymerase: Roles for DNA and protein determinants. Proceedings of the National Academy of Sciences 94, 9538–9543. 10.1073/pnas.94.18.9538.

15. Chaney, M., and Buck, M. (1999). The sigma 54 DNA-binding domain includes a determinant of enhancer responsiveness. Mol Microbiol 33, 1200–1209. 10.1046/j.1365-2958.1999.01566.x.

16. Campbell, E.A., Kamath, S., Rajashankar, K.R., Wu, M., and Darst, S.A. (2017). Crystal structure of Aquifex aeolicus σN bound to promoter DNA and the structure of σN-holoenzyme. Proceedings of the National Academy of Sciences 114, E1805–E1814. 10.1073/pnas.1619464114.

17. Merrick, M., and Chambers, S. (1992). The helix-turn-helix motif of sigma 54 is involved in recognition of the −13 promoter region. J Bacteriol 174, 7221–7226. 10.1128/jb.174.22.7221-7226.1992.

18. Taylor, M., Butler, R., Chambers, S., Casimiro, M., Badii, F., and Merrick, M. (1996). The RpoN-box motif of the RNA polymerase sigma factor σ ^N^ plays a role in promoter recognition. Mol Microbiol 22, 1045–1054. 10.1046/j.1365-2958.1996.01547.x.

19. Morris, L., Cannon, W., Claverie-Martin, F., Austin, S., and Buck, M. (1994). DNA distortion and nucleation of local DNA unwinding within sigma-54 (σN) holoenzyme closed promoter complexes. Journal of Biological Chemistry 269, 11563–11571. 10.1016/s0021-9258(19)78161-7.

20. Ye, F., Gao, F., Liu, X., Buck, M., and Zhang, X. (2022). Mechanisms of DNA opening revealed in AAA+ transcription complex structures. Sci Adv 8. 10.1126/sciadv.add3479.

21. Mueller, A.U., Chen, J., Wu, M., Chiu, C., Nixon, B.T., Campbell, E.A., and Darst, S.A. (2023). A general mechanism for transcription bubble nucleation in bacteria. Proceedings of the National Academy of Sciences 120. 10.1073/pnas.2220874120.

22. Chaney, M., Grande, R., Wigneshweraraj, S.R., Cannon, W., Casaz, P., Gallegos, M.-T., Schumacher, J., Jones, S., Elderkin, S., Dago, A.E., et al. (2001). Binding of transcriptional activators to sigma 54 in the presence of the transition state analog ADP-aluminum fluoride: insights into activator mechanochemical action. Genes Dev 15, 2282–2294. 10.1101/gad.205501.

23. Puchades, C., Sandate, C.R., and Lander, G.C. (2020). The molecular principles governing the activity and functional diversity of AAA+ proteins. Nat Rev Mol Cell Biol 21, 43–58. 10.1038/s41580-019-0183-6.

24. Stennett, E.M.S., Ciuba, M.A., Lin, S., and Levitus, M. (2015). Demystifying PIFE: The Photophysics Behind the Protein-Induced Fluorescence Enhancement Phenomenon in Cy3. J Phys Chem Lett 6, 1819–1823. 10.1021/acs.jpclett.5b00613.

25. Ko, J., and Heyduk, T. (2014). Kinetics of promoter escape by bacterial RNA polymerase: effects of promoter contacts and transcription bubble collapse. Biochemical Journal 463, 135–144. 10.1042/BJ20140179.

26. Studholme, D.J., Wigneshwereraraj, S.R., Gallegos, M.-T., and Buck, M. (2000). Functionality of Purified ς ^N^ (ς ^54^) and a NifA-Like Protein from the Hyperthermophile *Aquifex aeolicus*. J Bacteriol 182, 1616–1623. 10.1128/JB.182.6.1616-1623.2000.

27. Chen, B., Sysoeva, T.A., Chowdhury, S., Guo, L., and Nixon, B.T. (2009). ADPase activity of recombinantly expressed thermotolerant ATPases may be caused by copurification of adenylate kinase of Escherichia coli. FEBS Journal 276, 807–815. 10.1111/j.1742-4658.2008.06825.x.

28. Friedman, L.J., and Gelles, J. (2012). Mechanism of Transcription Initiation at an Activator-Dependent Promoter Defined by Single-Molecule Observation. Cell 148, 679–689. 10.1016/j.cell.2012.01.018.

29. Wang, J.T., and Gralla, J.D. (1996). The transcription initiation pathway of sigma 54 mutants that bypass the enhancer protein requirement: Implications for the mechanism of activation. Journal of Biological Chemistry 271, 32707–32713. 10.1074/jbc.271.51.32707.

30. Casaz, P., Gallegos, M.-T., and Buck, M. (1999). Systematic analysis of σ 54 N-terminal sequences identifies regions involved in positive and negative regulation of transcription. J Mol Biol 292, 229–239. 10.1006/jmbi.1999.3076.

31. Wang, L., and Gralla, J.D. (2001). Roles for the C-terminal Region of Sigma 54 in Transcriptional Silencing and DNA Binding. Journal of Biological Chemistry 276, 8979–8986. 10.1074/jbc.M009587200.

32. Kavalchuk, M., Jomaa, A., Müller, A.U., and Weber-Ban, E. (2022). Structural basis of prokaryotic ubiquitin-like protein engagement and translocation by the mycobacterial Mpa-proteasome complex. Nat Commun 13, 276. 10.1038/s41467-021-27787-3.

33. Cannon, W., Wigneshweraraj, S.R., and Buck, M. (2002). Interactions of regulated and deregulated forms of the sigma54 holoenzyme with heteroduplex promoter DNA. Nucleic Acids Res 30, 886–893. 10.1093/nar/30.4.886.

34. Gallegos, M.-T., and Buck, M. (2000). Sequences in σ 54 region I required for binding to early melted DNA and their involvement in sigma-DNA isomerisation. J Mol Biol 297, 849–859. 10.1006/jmbi.2000.3608.

35. Lin, G., Tsu, C., Dick, L., Zhou, X.K., and Nathan, C. (2008). Distinct Specificities of Mycobacterium tuberculosis and Mammalian Proteasomes for N-Acetyl Tripeptide Substrates. Journal of Biological Chemistry 283, 34423–34431. 10.1074/jbc.M805324200.

36. Lin, G., Hu, G., Tsu, C., Kunes, Y.Z., Li, H., Dick, L., Parsons, T., Li, P., Chen, Z., Zwickl, P., et al. (2006). *Mycobacterium tuberculosis prcBA* genes encode a gated proteasome with broad oligopeptide specificity. Mol Microbiol 59, 1405–1416. 10.1111/j.1365-2958.2005.05035.x.

37. Sasse-Dwight, S., and Gralla, J.D. (1990). Role of eukaryotic-type functional domains found in the prokaryotic enhancer receptor factor σ54. Cell 62, 945–954. 10.1016/0092-8674(90)90269-K.

38. Chen, B., Sysoeva, T.A., Chowdhury, S., Guo, L., De Carlo, S., Hanson, J.A., Yang, H., and Nixon, B.T. (2010). Engagement of Arginine Finger to ATP Triggers Large Conformational Changes in NtrC1 AAA+ ATPase for Remodeling Bacterial RNA Polymerase. Structure 18, 1420–1430. 10.1016/j.str.2010.08.018.

39. Nadanaciva, S., Weber, J., Wilke-Mounts, S., and Senior, A.E. (1999). Importance of F_1_ -ATPase Residue α-Arg-376 for Catalytic Transition State Stabilization. Biochemistry 38, 15493–15499. 10.1021/bi9917683.

40. Bae, B., Davis, E., Brown, D., Campbell, E.A., Wigneshweraraj, S., and Darst, S.A. (2013). Phage T7 Gp2 inhibition of *Escherichia coli* RNA polymerase involves misappropriation of σ ^70^ domain 1.1. Proceedings of the National Academy of Sciences 110, 19772–19777. 10.1073/pnas.1314576110.

41. Yang, Y., Darbari, V.C., Zhang, N., Lu, D., Glyde, R., Wang, Y.-P., Winkelman, J.T., Gourse, R.L., Murakami, K.S., Buck, M., et al. (2015). Structures of the RNA polymerase-54 reveal new and conserved regulatory strategies. Science (1979) 349, 882–885. 10.1126/science.aab1478.

42. Glyde, R., Ye, F., Jovanovic, M., Kotta-Loizou, I., Buck, M., and Zhang, X. (2018). Structures of Bacterial RNA Polymerase Complexes Reveal the Mechanism of DNA Loading and Transcription Initiation. Mol Cell 70, 1111–1120.e3. 10.1016/j.molcel.2018.05.021.

43. Kardon, J.R., Moroco, J.A., Engen, J.R., and Baker, T.A. (2020). Mitochondrial ClpX activates an essential biosynthetic enzyme through partial unfolding. Elife 9. 10.7554/eLife.54387.

44. Ye, Q., Rosenberg, S.C., Moeller, A., Speir, J.A., Su, T.Y., and Corbett, K.D. (2015). TRIP13 is a protein-remodeling AAA+ ATPase that catalyzes MAD2 conformation switching. Elife 4. 10.7554/eLife.07367.

45. Alfieri, C., Chang, L., and Barford, D. (2018). Mechanism for remodelling of the cell cycle checkpoint protein MAD2 by the ATPase TRIP13. Nature 559, 274–278. 10.1038/s41586-018-0281-1.

46. Bhat, J.Y., Miličić, G., Thieulin-Pardo, G., Bracher, A., Maxwell, A., Ciniawsky, S., Mueller-Cajar, O., Engen, J.R., Hartl, F.U., Wendler, P., et al. (2017). Mechanism of Enzyme Repair by the AAA+ Chaperone Rubisco Activase. Mol Cell 67, 744–756.e6. 10.1016/j.molcel.2017.07.004.

47. Ryu, J.-K., Min, D., Rah, S.-H., Kim, S.J., Park, Y., Kim, H., Hyeon, C., Kim, H.M., Jahn, R., and Yoon, T.-Y. (2015). Spring-loaded unraveling of a single SNARE complex by NSF in one round of ATP turnover. Science (1979) 347, 1485–1489. 10.1126/science.aaa5267.

48. White, K.I., Zhao, M., Choi, U.B., Pfuetzner, R.A., and Brunger, A.T. (2018). Structural principles of SNARE complex recognition by the AAA+ protein NSF. Elife 7. 10.7554/eLife.38888.

49. Burrows, P.C., Joly, N., Cannon, W. V., Cámara, B.P., Rappas, M., Zhang, X., Dawes, K., Nixon, B.T., Wigneshweraraj, S.R., and Buck, M. (2009). Coupling σ Factor Conformation to RNA Polymerase Reorganisation for DNA Melting. J Mol Biol 387, 306–319. 10.1016/j.jmb.2009.01.052.

50. Murakami, K.S., Masuda, S., and Darst, S.A. (2002). Structural Basis of Transcription Initiation: RNA Polymerase Holoenzyme at 4 Å Resolution. Science (1979) 296, 1280–1284. 10.1126/science.1069594.

51. Zhang, Y., Feng, Y., Chatterjee, S., Tuske, S., Ho, M.X., Arnold, E., and Ebright, R.H. (2012). Structural Basis of Transcription Initiation. Science (1979) 338, 1076–1080. 10.1126/science.1227786.

52. Kulbachinskiy, A., and Mustaev, A. (2006). Region 3.2 of the σ Subunit Contributes to the Binding of the 3ʹ-Initiating Nucleotide in the RNA Polymerase Active Center and Facilitates Promoter Clearance during Initiation. Journal of Biological Chemistry 281, 18273–18276. 10.1074/jbc.C600060200.

53. Campbell, E.A., Muzzin, O., Chlenov, M., Sun, J.L., Olson, C.A., Weinman, O., Trester-Zedlitz, M.L., and Darst, S.A. (2002). Structure of the Bacterial RNA Polymerase Promoter Specificity σ Subunit. Mol Cell 9, 527–539. 10.1016/S1097-2765(02)00470-7.

54. Gao, F., Ye, F., Zhang, B., Cronin, N., Buck, M., and Zhang, X. (2024). Structural basis of σ ^54^ displacement and promoter escape in bacterial transcription. Proceedings of the National Academy of Sciences 121. 10.1073/pnas.2309670120.

55. Chen, J., Gopalkrishnan, S., Chiu, C., Chen, A.Y., Campbell, E.A., Gourse, R.L., Ross, W., and Darst, S.A. (2019). E. coli TraR allosterically regulates transcription initiation by altering RNA polymerase conformation. Elife 8. 10.7554/eLife.49375.

56. Wulf, M.G., Maguire, S., Dai, N., Blondel, A., Posfai, D., Krishnan, K., Sun, Z., Guan, S., and Corrêa, I.R. (2022). Chemical capping improves template switching and enhances sequencing of small RNAs. Nucleic Acids Res 50, e2–e2. 10.1093/nar/gkab861.

57. Mastronarde, D.N. (2005). Automated electron microscope tomography using robust prediction of specimen movements. J Struct Biol 152, 36–51. 10.1016/j.jsb.2005.07.007.

58. Zheng, S.Q., Palovcak, E., Armache, J.-P., Verba, K.A., Cheng, Y., and Agard, D.A. (2017). MotionCor2: anisotropic correction of beam-induced motion for improved cryo-electron microscopy. Nat Methods 14, 331–332. 10.1038/nmeth.4193.

59. Punjani, A., Rubinstein, J.L., Fleet, D.J., and Brubaker, M.A. (2017). cryoSPARC: algorithms for rapid unsupervised cryo-EM structure determination. Nat Methods 14, 290–296. 10.1038/nmeth.4169.

60. Punjani, A., Zhang, H., and Fleet, D.J. (2020). Non-uniform refinement: adaptive regularization improves single-particle cryo-EM reconstruction. Nat Methods 17, 1214–1221. 10.1038/s41592-020-00990-8.

61. Zivanov, J., Nakane, T., Forsberg, B.O., Kimanius, D., Hagen, W.J., Lindahl, E., and Scheres, S.H. (2018). New tools for automated high-resolution cryo-EM structure determination in RELION-3. Elife 7. 10.7554/eLife.42166.

62. Liebschner, D., Afonine, P. V., Baker, M.L., Bunkóczi, G., Chen, V.B., Croll, T.I., Hintze, B., Hung, L.-W., Jain, S., McCoy, A.J., et al. (2019). Macromolecular structure determination using X-rays, neutrons and electrons: recent developments in Phenix. Acta Crystallogr D Struct Biol 75, 861–877. 10.1107/S2059798319011471.

63. Cardone, G., Heymann, J.B., and Steven, A.C. (2013). One number does not fit all: Mapping local variations in resolution in cryo-EM reconstructions. J Struct Biol 184, 226–236. 10.1016/j.jsb.2013.08.002.

64. Morin, A., Eisenbraun, B., Key, J., Sanschagrin, P.C., Timony, M.A., Ottaviano, M., and Sliz, P. (2013). Collaboration gets the most out of software. Elife 2. 10.7554/eLife.01456.

65. Sysoeva, T.A., Chowdhury, S., Guo, L., and Nixon, B.T. (2013). Nucleotide-induced asymmetry within ATPase activator ring drives 54-RNAP interaction and ATP hydrolysis. Genes Dev 27, 2500–2511. 10.1101/gad.229385.113.

66. Adams, P.D., Afonine, P. V., Bunkóczi, G., Chen, V.B., Davis, I.W., Echols, N., Headd, J.J., Hung, L.-W., Kapral, G.J., Grosse-Kunstleve, R.W., et al. (2010). PHENIX: a comprehensive Python-based system for macromolecular structure solution. Acta Crystallogr D Biol Crystallogr 66, 213–221. 10.1107/S0907444909052925.

67. Emsley, P., Lohkamp, B., Scott, W.G., and Cowtan, K. (2010). Features and development of Coot. Acta Crystallogr D Biol Crystallogr 66, 486–501. 10.1107/S0907444910007493.

68. Croll, T.I. (2018). *ISOLDE* : a physically realistic environment for model building into low-resolution electron-density maps. Acta Crystallogr D Struct Biol 74, 519–530. 10.1107/S2059798318002425.

69. Meng, E.C., Goddard, T.D., Pettersen, E.F., Couch, G.S., Pearson, Z.J., Morris, J.H., and Ferrin, T.E. (2023). UCSF ChimeraX : Tools for structure building and analysis. Protein Science 32. 10.1002/pro.4792.

70. Sievers, F., Wilm, A., Dineen, D., Gibson, T.J., Karplus, K., Li, W., Lopez, R., McWilliam, H., Remmert, M., Söding, J., et al. (2011). Fast, scalable generation of high-quality protein multiple sequence alignments using Clustal Omega. Mol Syst Biol 7, 539. 10.1038/msb.2011.75.

71. Crooks, G.E., Hon, G., Chandonia, J.-M., and Brenner, S.E. (2004). WebLogo: A Sequence Logo Generator. Genome Res 14, 1188–1190. 10.1101/gr.849004.

