## Supplementary information for "Real-time capture of σ^N^ transcription initiation intermediates reveals mechanism of ATPase-driven activation by limited unfolding"

The authors declare no conflict of interest

Supplementary information includes 4 tables.

**Supplementary Table 1: 5'-RACE/template switching results establish transcription start site (TSS) from linear templates dhsU and dhsU+2T.** The sequence column shows the aligned sequences in FASTA format. The adapter sequence is marked in bold. To assign the 5'-end, either 3 G (for uncapped RNAs) or 4 G (for capped RNAs) of the adapter were removed (see methods for details). Bases in parentheses indicate ambiguity of 5'-end assignment (cap or TSS).

| Template | Aligned RACE sequence* | Assigned 5' end |
|---|---|---|
| dhsU | **GTACATGGG**GGAACAAAAATGGAGGTAAGAGTATGGGTGG | 5'-GAACAA |
| dhsU | **GTACATGGG**GGAACAAAAATGGAGGTAAGAGTATGGGTGG | 5'-GAACAA |
| dhsU | **GTACATGGG**GA-ACAAAAATGGAGGTAAGAGTATGGGTGG | 5'-(G)AACAA |
| dhsU | **GTACATGGG**GGAACAAAAATGGAGGTAAGAGTATGGGTGG | 5'-GAACAA |
| dhsU | **GTACATGGG**GGAACAAAAATGGAGGTAAGAGTATGGGTGG | 5'-GAACAA |
| dhsU | **GTACATGGG**GA-ACAAAAATGGAGGTAAGAGTATGGGT-G | 5'-(G)AACAA |
| dhsU | **GTACATGGG**GGAACAAAAATGGAGGTAAGAGTATGGGTGG | 5'-GAACAA |
| dhsU | **GTACATGGG**GA--CAAAAATGGAGGTAAGAGTATGGGTGG | 5'-ACAAAA |
| dhsU | **GTACATGGG**GA-ACAAAAATGGAGGTAAGAGTATGGGTGG | 5'-GAACAA |
| dhsU+2T | **GTACATGGG**GGTACAAAAATGGAGGTAAGAGTATGGGTGG | 5'-GTACAA |
| dhsU+2T | **GTACATGGG**GAA------ATGGAGGTAAGAGTATGGGTGG | 5'-AAATGG |
| dhsU+2T | **GTACATGGG**GGTACAAAAATGGAGGTAAGAGTATG-GTGG | 5'-GTACAA |
| dhsU+2T | **GTACATGGG**GTA-CAAAAATGGAGGTAAGAGTATGGGTGG | 5'-GTACAA |
| dhsU+2T | **GTACATGGG**GT-ACAAAAATGGAGGTAAGAGTATGGGTGG | 5'-GTACAA |
| dhsU+2T | **GTACATGGG**GGTACAAAAATGGAGGTAAGAGTATGGGTGG | 5'-GTACAA |
| dhsU+2T | **GTACATGGG**GGTACAAAAATGGAGGTAAAATTATGGGTGG | 5'-GTACAA |

* Total of 10 sequencing reactions per template: removed 1 failed reaction (dhsU+2T) and 3 special cases (end binding/template jump; 1 for dhsU, 2 for dhsU+2T) before alignment.

**Supplementary Table 2: Oligonucleotides.** Complementary overhangs in lowercase letters.

| Name | Sequence (5’-> 3’) | Remarks |
|---|---|---|
| aaeC1-GS4GQYL_fw | ctggggcagtatctctaatgaGGATCCGAATTCGAGCTCCG | |
| aaeC1-GS4GQYL_rv | ggagccggacccTTTGCTATTAACCAGACAGCTCAGTTC | |
| dhsU_top | CGCAAGTTCCTTAGAATTTCAGTGTCCAGAAATTGGCACGAAAATTGCAATAAATACAACGAACAAAAATGGAGGTAAGAGTATGGGTGG | Promoter fragment |
| dhsU_bot | CCACCCATACTCTTACCTCCATTTTTGTTCGTTGTATTTATTGCAATTTTCGTGCCAATTTCTGGACACTGAAATTCTAAGGAACTTGCG | Promoter fragment |
| dhsU-CT_top | CGCAAGTTCCTTAGAATTTCAGTGTCCAGAAATTGGCACGAAAATTGCCTTAAATACAACGAACAAAAATGGAGGTAAGAGTATGGGTGG | pre-melted 2-nt bubble; non-complementary bases underlined |
| dhsU+2T_top | CGCAAGTTCCTTAGAATTTCAGTGTCCAGAAATTGGCACGAAAATTGCAATAAATACAACGTACAAAAATGGAGGTAAGAGTATGGGTGG | Promoter fragment |
| dhsU+2T_bot | CCACCCATACTCTTACCTCCATTTTTGTACGTTGTATTTATTGCAATTTTCGTGCCAATTTCTGGACACTGAAATTCTAAGGAACTTGCG | Promoter fragment |
| dhsU-RTprim | GAGATTCGTATGCCTGGTATGGATCGCTTGGCAccacccatactcttacctc | Gene-specific part in lowercase; adapter sequence for PCR primers underlined |
| RACE-TSO_fw | GCTAATCATTGCAAGCAGTGGTATCAACGCAGAGTACATrGrGrG | rG = riboguanosine; adapter sequence for PCR primer underlined |
| RACE-PCR-TSO_fw | CATTGCAAGCAGTGGTATCAAC | cDNA amplification |
| RACE-PCR-RT_rv | CGTATGCCTGGTATGGATCG | cDNA amplification |
| RACE-pTwist_fw | gatccataccaggcatacgAGGCTAGGTGGAGGCTC | Vector linearization |
| RACE-pTwist_rv | gataccactgcttgcaatgAGGTCAGGCGGAATGGC | Vector linearization |
| M13-40FOR | GTTTTCCCAGTCACGAC | Colony sequencing |
| dhsU+2T-Cy3_top | CGCAAGTTCCTTAGAATTTCAGTGTCCAGAAATTGGCACGAAAATTGCAATAAATACAACG/iCy3N/ACAAAAATGGAGGTAAGAGTATGGGTGG | iCy3N = Cy3-NHS ester modification on deoxythymidine |

**Supplementary Table 3: Results of the N-terminal protein sequencing by Edman degradation.** Transcription complexes with the indicated $\sigma^N$ variants were formed on dhsU-CT in presence of C1-GS4 and 20S-og in absence or presence of ATP. Parentheses contain minor peaks and/or less confident calls. n/d = not determined. X = any residue/no residue assignment was possible.

| Cycle | $\sigma^N$ WT | $\sigma^N$ WT + ATP | Nt-$\sigma^N$ | Nt-$\sigma^N$ + ATP | GS7-$\sigma^N$ | GS7-$\sigma^N$ + ATP |
|---|---|---|---|---|---|---|
| 1 | M | M, (T) | D, (M, S, T, K) | L, (M, S) | M | L, (M) |
| 2 | K | K, (I, P, V, L, G, A) | I, (K, G, Y) | A, (K) | K | A, (K) |
| 3 | Q | Q, (E, Y, S, F) | Q, (G, V) | M | Q | M |
| 4 | G | G, (L, T) | G, (F) | K, (T, G) | G | G, (K) |
| 5 | L | L, (E, T) | L | Q, (P, L) | L | S, (P, L) |
| 6 | n/d | Q, (S) | n/d | G | n/d | G, (Q) |
| 7 | n/d | L, (A, D) | n/d | L, (A) | n/d | S |
| 8 | n/d | R, (I, D) | n/d | Q | n/d | X |
| 9 | n/d | L, (Y) | n/d | L | n/d | X |
| 10 | n/d | S, (G) | n/d | R, (D) | n/d | (G, A) |
| 11 | n/d | Q | n/d | (L, I) | n/d | (M, I) |
| 12 | n/d | Q | n/d | (S) | n/d | (K, E) |
| 13 | n/d | L | n/d | (Q) | n/d | (Q, L) |
| 14 | n/d | A, (D) | n/d | (Q, A) | n/d | (G, A) |
| 15 | n/d | M | n/d | (L) | n/d | (L) |

**Supplementary Table 4: Parameter estimates from the kinetic data fitting.** Values for 95% confidence interval are given in parentheses.

| Dataset | a0 | a1 | k |
|---|---|---|---|
| 50 nM | 0.0145<br>(0.014 to 0.015) | 0.3212<br>(0.3199 to 0.3266) | 0.0066<br>(0.0065 to 0.0067) |
| 100 nM | −0.0059<br>(−0.0077 to −0.0040) | 1.018<br>(1.014 to 1.023) | 0.0086<br>(0.0085 to 0.0087) |
| 200 nM | −0.0049<br>(−0.0075 to −0.0092) | 1.345<br>(1.339 to 1.351) | 0.0091<br>(0.0090 to 0.0092) |
